# Microtubule disassembly by caspases is the rate-limiting step of cell extrusion

**DOI:** 10.1101/2021.10.15.464503

**Authors:** Alexis Villars, Alexis Matamoro-Vidal, Florence Levillayer, Romain Levayer

## Abstract

Epithelial cell death is essential for tissue homeostasis, robustness and morphogenesis. The expulsion of epithelial cells following caspase activation requires well-orchestrated remodeling steps leading to cell elimination without impairing tissue sealing. While numerous studies have provided insight about the process of cell extrusion, we still know very little about the relationship between caspase activation and the remodeling steps of cell extrusion. Moreover, most studies of cell extrusion focused on the regulation of actomyosin and steps leading to the formation of a supracellular contractile ring. However, the contribution of other cellular factors to cell extrusion has been poorly explored. Using the *Drosophila* pupal notum, a single layer epithelium where most extrusion events are caspase-dependent, we first showed that the initiation of cell extrusion and apical constriction are surprisingly not associated with the modulation of actomyosin concentration/dynamics. Instead, cell apical constriction is initiated by the disassembly of a medio-apical mesh of microtubules which is driven by effector caspases. We confirmed that local and rapid increase/decrease of microtubules is sufficient to respectively expand/constrict cell apical area. Importantly, the depletion of microtubules is sufficient to bypass the requirement of caspases for cell extrusion. This study shows that microtubules disassembly by caspases is a key rate-limiting steps of extrusion, and outlines a more general function of microtubules in epithelial cell shape stabilisation.

## Introduction

How epithelia maintain their physical and chemical barrier functions despite their inherent dynamics due to cell proliferation and cell death is a central question of epithelial biology. Cell extrusion, a sequence of coordinated remodeling steps leading to cell expulsion, is an essential process to conciliate high rates of apoptosis/cell elimination while preserving tissue sealing^1, 2^ This process is essential for tissue homoeostasis and its perturbation can lead to chronic inflammation or contribute to tumoural cell dissemination^2, 3^ Yet much remains unknown about extrusion regulation and orchestration.

Studies in the last decade have demonstrated that the remodelling steps of extrusions are mainly dependent on actomyosin contraction and mechanical coupling through E-cadherin (E-cad) adhesion. First, an actomyosin ring forms in the extruding cell driving cell-autonomous constriction^4–6^. This ring pulls on neighbouring cells through E-cad anchorage, resulting in force transmission which promotes the recruitment of actomyosin in the neighbouring cells and the formation of a supracellular actomyosin cable^1, 4–7^ Eventually the constriction of the cable combined with E-cad disassembly^6,8^ lead to cell expulsion either on the apical or the basal side of the tissue. Meanwhile, neighbouring basal protrusions also contribute to cell detachment^9,10^ Alternatively, pulses of contractile medio-apical actomyosin can also contribute to cell expulsion^11,12^ Interestingly, while a lot of emphasis has been given to actomyosin and E-cad regulation, we only have a limited understanding of the contribution of other cellular factors/cytoskeleton components to cell extrusion (see^13^ for one exception). Apoptosis is one of the main mode of programmed cell death which is essential for tissue homeostasis and morphogenesis^14^. It is driven by the activation of caspases which through the cleavage of thousands of proteins orchestrate cell deconstruction^15^. While caspase activation is an important mode of epithelial cell elimination^14^, how caspases orchestrate the key steps of extrusion remain poorly understood. Accordingly, only a handful of caspases targets relevant for cell extrusion has been identified so far^16^.

The morphogenesis of the *Drosophila* pupal notum, a single layer epithelium located in the back of the thorax, is an ideal system to study the regulation of apoptosis and cell extrusion. High rates of cell extrusion in reproducible patterns are observed in the midline and posterior region of the notum^17–21^. Interestingly, the majority of cell extrusion events in the pupal notum are effector caspase-dependent. Accordingly, caspase activation always precedes cell extrusion and inhibition of caspase in clones or throughout the tissue dramatically reduces the rate of cell extrusion^18,20–22^. However, we currently do not know which steps of extrusion are regulated by caspases or how effector caspase activation initiates and orchestrates epithelial cell extrusion.

Here, we first performed the quantitative phenomenology of cell extrusion in the midline of the pupal notum. Surprisingly, while we observed the formation of a supracellular actomyosin ring in the late phase of extrusion, the initiation of cell apical constriction was not associated with any change in the dynamics/concentration of actomyosin and Rho. Accordingly, comparison with the behaviour of extruding cells in a vertex model suggested that cell extrusion in the notum is not initiated by a change of line tension. Instead, we found that cell extrusion initiation is concomitant with the disassembly of an apical mesh of microtubules (MTs). This disassembly is effector caspase-dependent and is required for cell extrusion. More importantly, the requirement of caspase activation for extrusion can be bypassed by MT disassembly, suggesting that the remodelling of MTs by caspases is an important rate-limiting step of cell extrusion. This work also emphasises the need to study the contribution of microtubules to epithelial cell shape regulation independently of actomyosin regulation.

## Results

### Actomyosin modulation is not responsible for the initiation of cell extrusion

We focused on the *Drosophila* pupal notum midline (**Figure 1A, B**), a region showing high rates of cell death and cell extrusion^17–20^. To better characterise the process of cell extrusion, we first quantified the evolution over time of the main regulators of cellcell adhesion (E-cad) and cortical tension (Non-muscle Myosinll, Myoll) by averaging and temporally aligning several extruding cells (one extrusion event lasting approximatively 30 min). Contrary to other tissues^6, 8^, we did not observe a depletion of E-cad at the junctions during the constriction process, but rather a progressive increase of its concentration (**Figure 1C, D, movie S1**). More strikingly, the onset of cell extrusion (defined by the inflexion of the apical perimeter, see **Methods**) was not associated with a clear increase of MyoII levels (looking at Myosin Regulatory Light Chain, MRLC, **Figure 1E, F, movie S2**), either at the junctional pool or in the medio-apical region (**Figure S1A**). Instead, a clear accumulation of Myoll forming a supracellular cable was observed during the last 10 minutes of cell extrusion (**Figure 1E, F, Figure S1A**). Accordingly, F-actin starts to accumulate only in the late phase of extrusion concomitantly with the formation of the supracellular cable (**Figure 1G,H, Figure S1G,H, movie S3**), similar to the dynamics we observed for Rho1, a central regulator of F-actin, MyoII activity and pulsatility^23^ (**Figure S1 I-J’, movie S4**).

**Figure 1:**
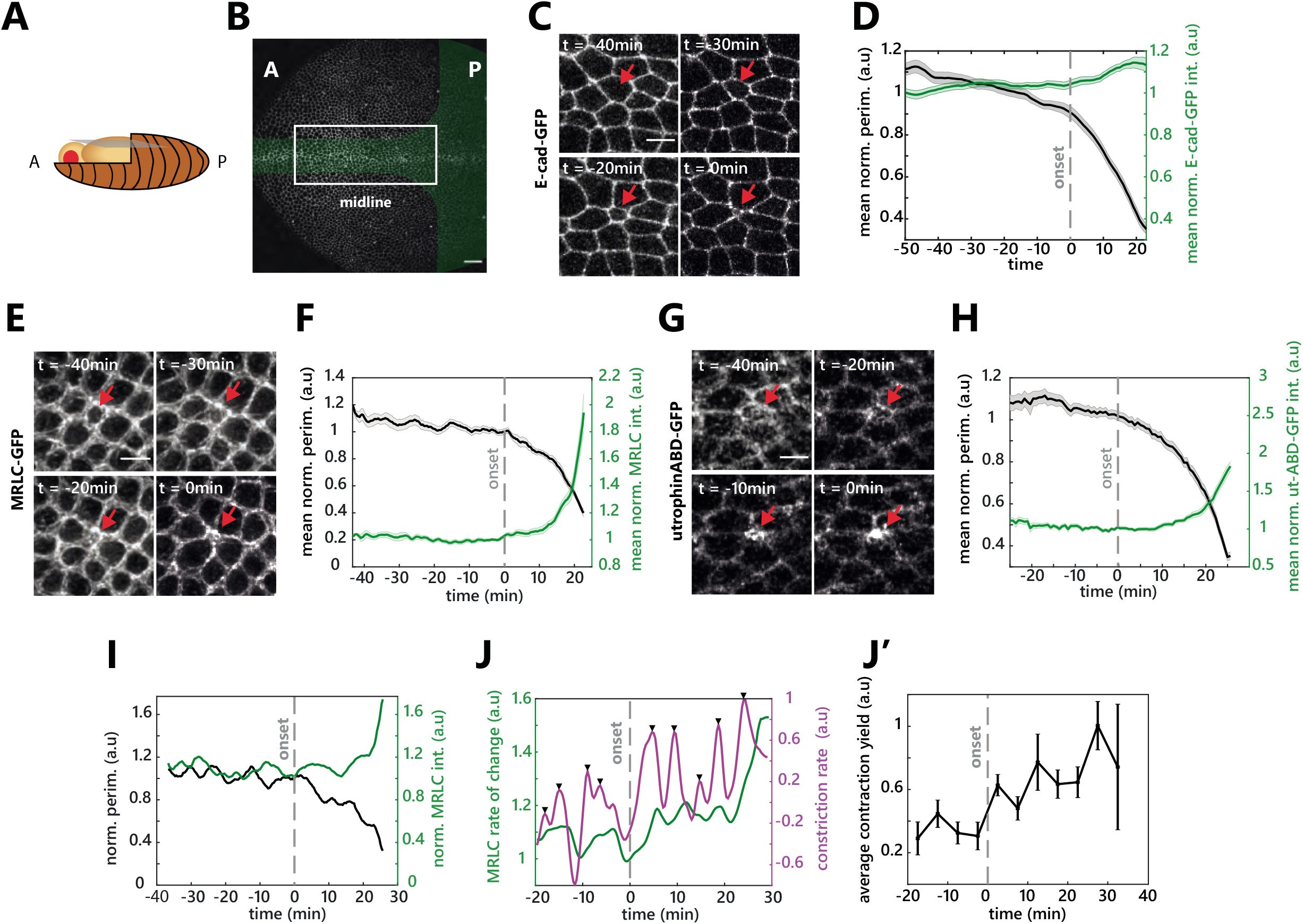
Actomyosin modulation is not responsible for the initiation of cell extrusion. **A:** Schematic of a *Drosophila* pupae. Orientation (used for every figure), anterior on the left and posterior on the right. The blue plane shows the notum. **B:** Notum at 16h After Pupal Formation (APF). The green zone represents the area with a high rate of cell elimination. White rectangle highlights the midline region used for extrusion analysis. A: Anterior, P: Posterior. Scale bar, 25μm. **C:** Snapshots of E-cad-GFP during cell extrusion. Red arrow shows an extruding cell. t0 is the time of extrusion termination (no apical area visible). Scale bar, 5μm. **D:** Averaged and normalised E-cad-GFP junctional signal during cell extrusion (green) and averaged normalised cell perimeter (black), light colour areas are s.e.m.. The curves were aligned temporally by using the extrusion termination time point (see **Methods**). Grey dotted line represents the onset of extrusion marked by the inflection of the perimeter curve (see **Methods**). N=2 pupae, n=27 cells. **E:** Snapshots of sqh-GFP (MRLC) during cell extrusion. Red arrow shows an extruding cell. t0 is the time of extrusion termination. Scale bar, 5μm. **F:** Averaged normalised sqh-GFP (MRLC) total signal (medial+junctional) during cell extrusion (green) and average normalised cell perimeter (black). Grey dotted line shows the onset of extrusion, light colour areas are s.e.m.. N=2 pupae, n=15 cells. **G:** Snapshots of actin during cell extrusion (visualized with utrophin Actin-Binding domain fused to GFP – utABD-GFP). Red arrow points at an extruding cell. t0 is the time of extrusion termination. Scale bar, 5μm. **H:** Averaged normalised utABD-GFP total signal (medial+junctional) during cell extrusion (green) and averaged normalised cell perimeter (black). Grey dotted line represents the onset of extrusion, light colour areas are s.e.m.. N=2 pupae, n=37 cells. **I.** Single cell representative curve of sqh-GFP (MRLC) total signal (green) and perimeter (black) showing MRLC pulsatility and perimeter fluctuations before and during cell extrusion. Grey dotted line represents the onset of extrusion. **J.** Single cell representative curve of sqh-GFP (MRLC) intensity rate of change (i.e. derivative, green) and the perimeter constriction rate (derivative of the perimeter, magenta). Black arrows show contraction pulses. Grey dotted line represents the onset of extrusion. **J’.** Averaged contraction yield (ratio of the constriction rate over MRCL junctional intensity) calculated in 5 min time windows (see **Methods**). Dotted line, extrusion onset, errors bars are s.e.m.. N=2 pupae, n=15 cells.

Pulsatile actomyosin recruitment is observed during a wide variety of morphogenetic processes^24–26^. We also observed fluctuating levels of MyoII (**Figure 1I, Figure S1B**) with pulses correlating with transient constriction of the cell apical perimeter (**Figure S1B, C**). The amplitude, duration and/or frequency of MyoII pulses can affect the efficiency of cell constriction^27^. However, we did not observe any significant change of these parameters before and after the onset of cell extrusion (**Figure S1D-F**). Finally, to better characterise the link between perimeter constriction and MyoII dynamics, we calculated a contraction yield (the ratio of constriction rate over the intensity of MyoII). We observed a significant increase in the contraction yield at the onset of cell extrusion (**Figure 1 J,J’**), suggesting that similar MyoII pulses lead to more deformation after the onset of extrusion.

Altogether, we found that the initiation of cell extrusion and apical constriction is not associated with a significant change of actin, MyoII and Rho dynamics/levels. Their enrichment appears during the last 10 minutes of extrusion and is associated with the formation of a supracellular actomyosin cable. This suggests that MyoII activation/recruitment and dynamics are not sufficient to explain the initiation of extrusion and that MyoII activation is unlikely to be the rate-limiting step that initiates cell extrusion. Accordingly, we observed a significant increase of MyoII levels upon inhibition of caspase activity (by depleting Hid, a proapoptotic gene, **Figure S1H, H’**), a condition that almost completely abolishes cell extrusion^20^, suggesting once again that MyoII recruitment is not the main rate limiting step of extrusion downstream of caspases.

### Cell extrusion in the midline is different from purse-string driven extrusion

To get a better understanding of the mechanical parameters regulating the initiation of cell extrusion in the midline, we used a 2D vertex model. The apical area of cells in the model can be modulated by two main parameters: the line tension, a by-product of junctional actomyosin and cell-cell adhesion which tend to minimise the perimeter of the cell, and the area elasticity, which constrains the variation in apical area of the cells and is thought to emerge from the incompressibility of cell volume and the properties of the medio-apical cortex^28,29^ The formation of a supracellular actomyosin cable, as observed in other instances of extrusion^1, 6^, should lead to an increase of line tension/contractility. We simulated such extrusion by implementing a progressive increase of the contractility parameter in a single cell (see **Methods**). This led to a progressive decrease in cell apical area concomitant with cell rounding (**Figure 2 A,C,E,G**, **movie S5** progressive increase of cell circularity), in good agreement with the profile of extrusion observed in the larval accessory cells of the abdomen which are driven by an actomyosin purse-string^6^ (**Figure 2 I**, **Figure S2A**). This however did not fit the pattern we observed in the midline of the pupal notum, where the initial phase of cell constriction is not associated with cell rounding (**Figure 2 J**). Cell constriction could also be initiated by a reduction of the cell resting area, which corresponds to the targeted apical area of cells in the absence of external constrains. Accordingly, the reduction of the resting area of a single cell in the vertex model led to cell constriction concomitant with a progressive reduction of cell roundness (**Figure 2 A,D,F,H**, **movie S5**), in good agreement with the dynamics we observed during the first 20 minutes of extrusion in the notum (**Figure 2J**). Altogether, this confirmed that the initiation of cell extrusion in the midline is unlikely to be driven by an increase in line tension/junction contractility, but rather by a process modulating cell resting area/compressibility.

**Figure 2:**
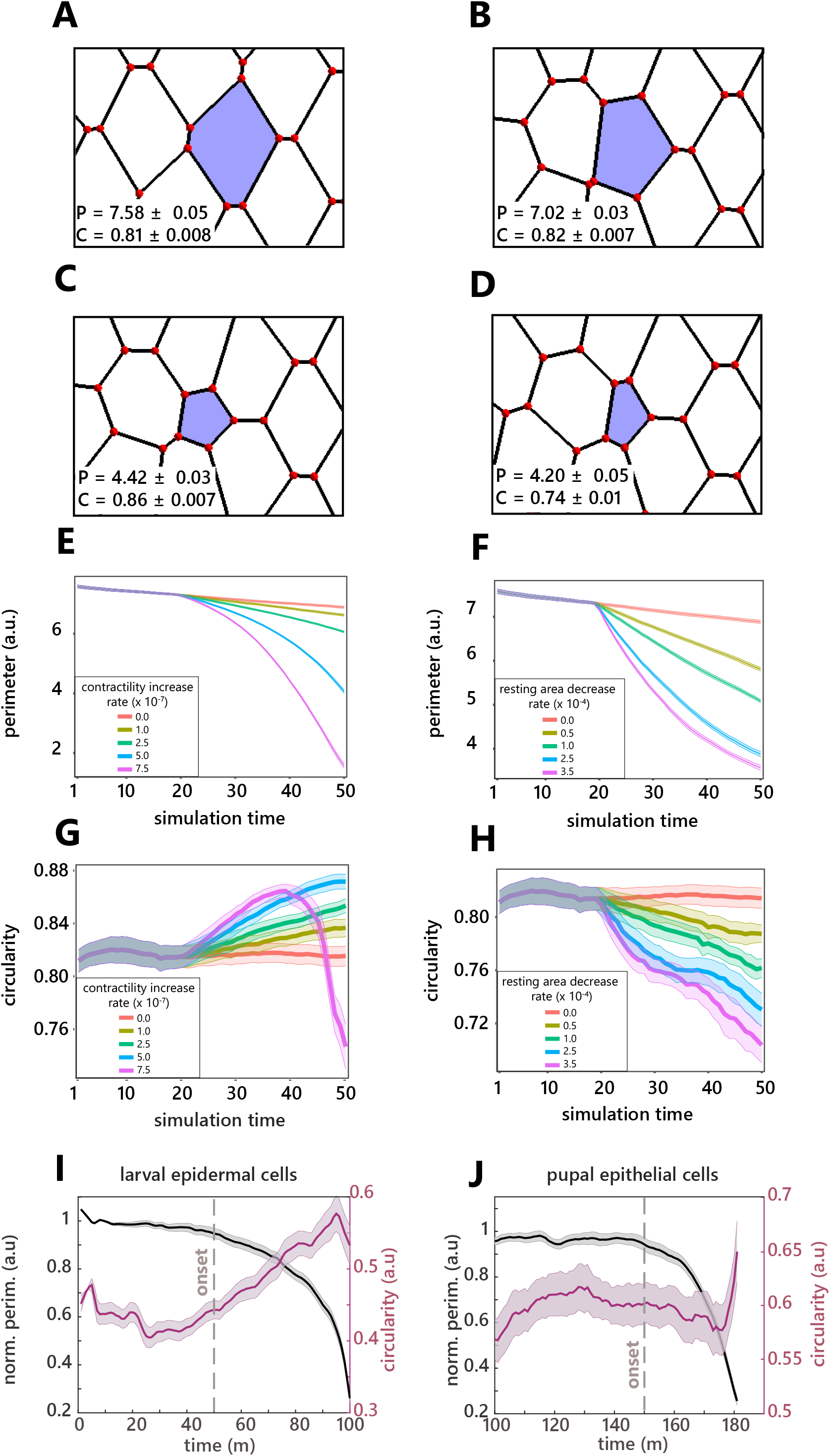
Cell extrusion in the midline is different from purse-string driven extrusion. **A-D:** Examples of a tracked cell (blue) from a vertex model during early extrusion phase at different simulation time steps (sts) and under different conditions. P is average perimeter ± s.e.m., and C denotes average circularity ± s.e.m. for the 10 tracked cells of the simulations. **A:** Initial state at sts = 1. **B:** Control simulation (no change of parameter in the blue cell) at sts = 40. **C:** Extrusion driven by increased cell contractility 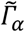 (contractility increase rate *c* = 7.5.10^-7^), at sts = 40 D: Extrusion through decreased resting area A_α_^(0)^ (resting area decrease rate *r* = 3.5.10^-4^) at sts = 40. **E-F**: Averaged cell perimeter ± s.e.m. for 10 tracked cells as a function of contractility increase rate (**E**) and of resting area decrease rate (**F**). Variation of contractility and resting area was initiated at sts = 20. **G-H**: Average cell circularity ± s.e.m. for 10 tracked cells as a function of contractility increase rate (**G**) and of resting area decrease rate (**H**). Variation of contractility and resting area was initiated at sts = 20. **I-J**: Averaged cell circularity (magenta) and averaged and normalised cell apical perimeter (black) during cell extrusion in larval epidermal cells, n= 37 cells, N = 2 pupae. (**I**) and in the pupal notum epithelial cells, n= 22 cells, N = 2 pupae (**J**). Light colour areas are s.e.m.. Grey dotted lines show the onset of extrusion.

### The disassembly of microtubules correlates with the onset of apical constriction

We next sought to identify which alternative factors could initiate cell extrusion in the midline of the pupal notum. Caspase activity can lead to a reduction of cell volume^30^, which could be responsible for the reduction of cell apical area. However, we did not observe a significant change in cell volume during the process of cell extrusion (**Figure S2B, C, movie S6**). Alternatively, a downregulation of extracellular matrix (ECM) binding on the cuticle side could facilitate apical area constriction. Yet we did not observe a modulation of integrin adhesion components at the onset of extrusion (**Figure S2D, movie S7**). We therefore checked the distribution of microtubules (MTs), which can also regulate epithelial cell shape^31, 32^ MT filaments accumulate in the medio-apical region of the midline cells as well as along the apico-basal axis (**Figure 3A, B, B’**). Strikingly, we observed a significant and reproducible depletion of apical MTs at the onset of cell extrusion (visualised with the MT-associated protein Jupiter-GFP, **Figure 3C-E, movie S8**). This depletion is concomitant with the onset of apical constriction (**Figure 3D, E**, peak of cross-correlation between MTs intensity and cell perimeter at t=2min). The same downregulation was observed with EB1-GFP, a marker of MT plus ends (**Figure S3 A-C, movie S9**), or upon expression of a tagged human α-tubulin (**Figure S3D, movie S10**). Interestingly, MT depletion in the extruding cell was followed later-on by an accumulation in the neighbouring cells of MTs close to the junctions shared with the dying cell (**Figure 3C and F-G’**, **Figure S3A-B, movies S8-S10**). This accumulation matches the timing of the actomyosin ring formation (**Figure 1E-H**) and is reminiscent of the MTs reorganisation previously described near MDCK extruding cells^33^. The loss of apico-medial MTs may be driven by the reorganisation of the non-centrosomal MTs regulators Patronin and Shot ^32, 34–37^. However we could not observe a clear modulation of their localisation/levels at the onset of cell extrusion (**Figure S3 E, F, movies S11-S12**). Alternatively, the apical disappearance of MTs may be driven by a basal shift of centrosomes (as observed in the dorsal folds of the early *Drosophila* embryo^32^), but we did not observe any change in the apico-basal position of the centrosomes at the onset of extrusion (**Figure S3G**). Finally, MTs downregulation may be driven by a global disassembly of MT filaments. Accordingly, the disappearance of MT filaments is not restricted to the most apical domain and seems to occur throughout the cell (**Figure S3H-I’**). Altogether, we observed a global disassembly of the MT network which is perfectly concomitant with the onset of cell extrusion and apical area reduction.

**Figure 3:**
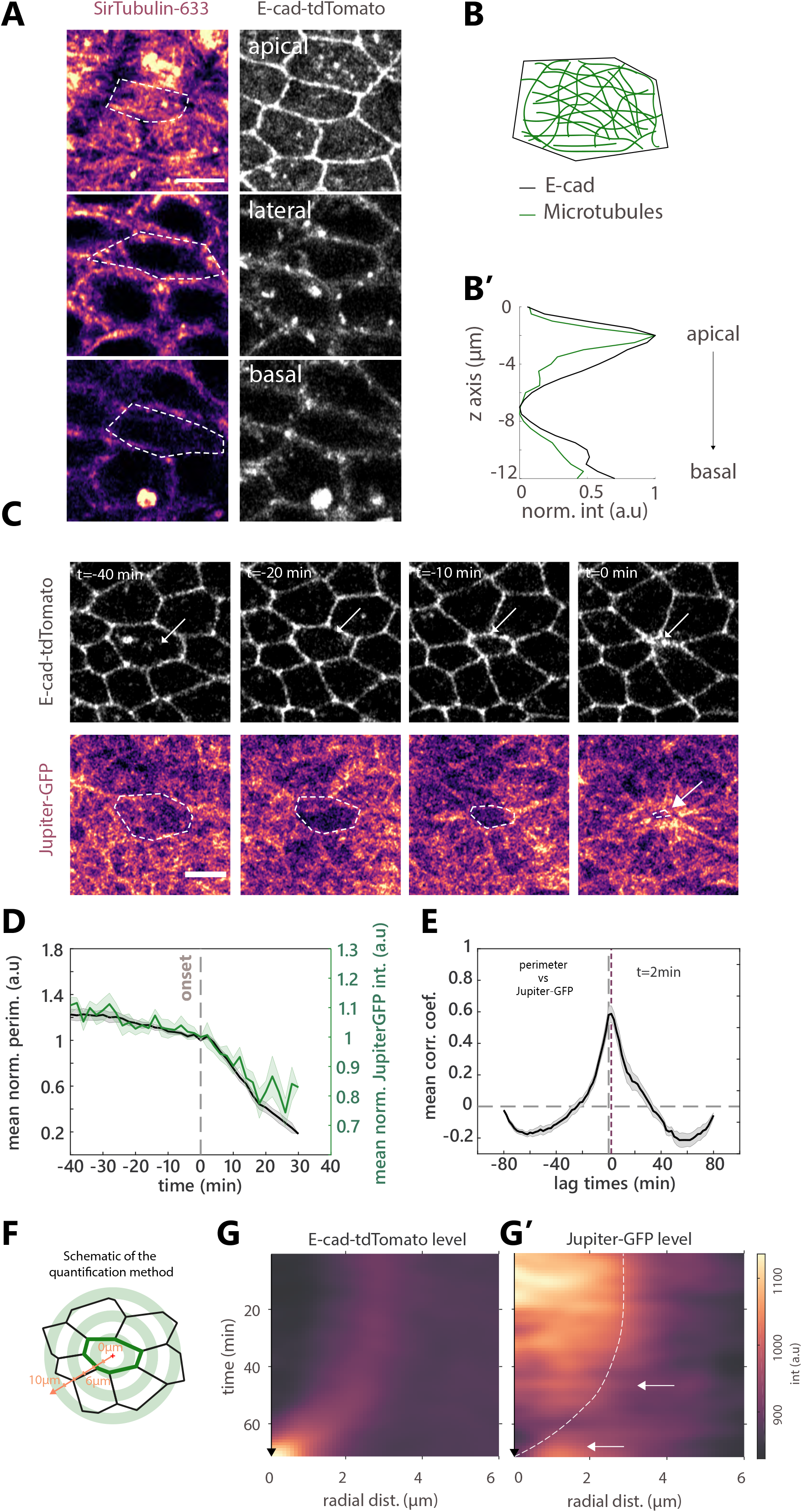
Microtubule depletion correlates with the onset of cell extrusion in the notum. **A.** Microtubules (MTs) visualised with sirTubulin (left, pseudo-colour) and E-cad-tdTomato (right, greyscale) along the apico-basal axis in a midline cell. White dotted lines show cell contour. Scale bar, 5μm. **B:** Schematic representation of MTs orientation in the junctional plane. **B’.** sirTubulin (green) and E-cad-tdTomato (black) total intensity along the apico-basal axis. 0 μm is the most apical plane. **C:** Snapshots of Jupiter-GFP (total tubulin pool, bottom, pseudo-colour) and E-cad-tdTomato (top, greyscale) during cell extrusion in the midline, white arrows and white dotted lines show the extruding cell. t0 min, termination of extrusion. Scale bar, 5μm. **D:** Averaged normalised Jupiter-GFP medial signal (green) and averaged normalized cell perimeter (black) during cell extrusion. Grey dotted line, extrusion onset, light colour areas, s.e.m.. t0 is the onset of extrusion. N = 2 pupae, n = 24 cells. **E:** Averaged normalised cross correlation of the cell perimeter vs Jupiter-GFP. Purple dotted line is at the maximum of correlation coefficient (t=2min). Horizontal grey dotted line is at correlation coefficient=0. Vertical dotted line is at lag time = 0 min. Light area is s.e.m.. n = 24 cells. F: Schematic of the method used to represent averaged MTs intensity profile in space and time during extrusion. Red cross shows the center of the extruding cell (junctions highlighted in green). The signal is measured every pixel in concentric bands of 3px thickness expanding from the cell center and the profile is averaged on every extruding cell analysed. **G:** Radial averaged kymograph (see **F**) of E-cad-tdTomato (left, pseudo-colour), time is on the y-axis going downward, x-axis is radial distance from the cell center. **G’**: Radial averaged kymograph of Jupiter-GFP signal. White dotted line represents the average cell contour (detected with the maximum of E-cad average signal for each time point). Top white arrow points at the onset of Jupiter-GFP depletion. Bottom arrow shows Jupiter-GFP accumulation in the neighbouring cells at the end of extrusion. n = 24 cells.

### MT depletion is effector caspase dependent

We then checked what could be responsible for MT disappearance. MT buckling driven by cell deformation can trigger MT disassembly^38–40^. As such, MT depletion during extrusion may be a cause or a consequence of the initiation of cell apical area constriction. To check if area constriction is sufficient to trigger MTs disassembly, we released tissue prestress in the notum by laser cutting a large tissue square^20, 41^. The transient constriction of cell apical area in the square region correlated with a transient increase of apical MT concentration (**Figure S4A-B, movie S13**), which is then followed by cell apical area re-expansion and MT intensity diminution. This suggests that cell apical constriction is not sufficient to disassemble MTs and that an active process must drive MT disassembly at the onset of extrusion. The activation of effector caspases is necessary for extrusion and always precedes cell constriction in the pupal notum^18, 20, 22^ We therefore checked whether effector caspase activation was necessary and sufficient for MT depletion. Previously, we developed an optogenetic tool (optoDronc ^21^) which can activate *Drosophila* Caspase9 triggering apoptosis and cell extrusion upon blue light exposure (**Figure 4B**). We observed a rapid depletion of MTs (visualised with injected sirTubulin) upon activation of optoDronc in clones which was concomitant with cell apical constriction (**Figure 4A,C, movie S14 top**). Activating optoDronc while inhibiting effector Caspases through p35 dramatically slowed-down the rate of cell extrusion (^21^ and **Figure 4D,E, movie S14 bottom**). While we could still see a late accumulation of MyoII in these slow constricting cells (**Figure S4C,** 3 hours post optoDronc activation), we could no longer observe MT depletion but rather a progressive increase in the apical concentration of MTs (**Figure 4D,E, movie S14 bottom**). This is in agreement with the numerous cells with low apical area and strong tubulin accumulation we observed in *hid-RMAi*/caspase-inhibited clones (**Figure 4G**). Interestingly, contrary to normal extrusions in the notum (**Figure 2J**), circularity progressively increases during perimeter constriction upon optoDronc activation in cells expressing p35 (**Figure 4F**). This fits with a slow constriction driven by an increase of contractility/line tension. Altogether, this suggests that MT depletion during extrusion is driven directly or indirectly by effector caspases activation.

**Figure 4:**
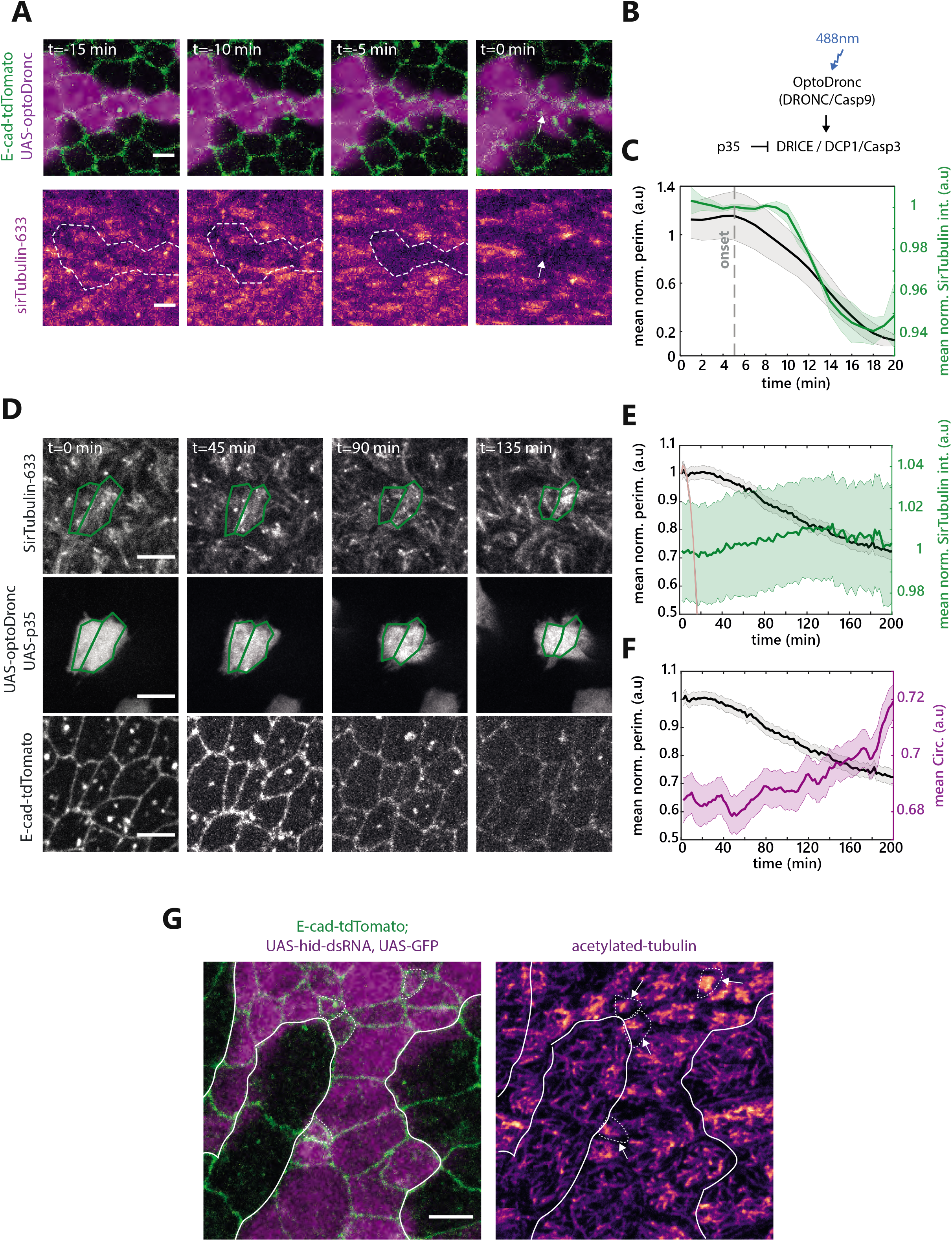
Microtubule depletion is effector caspase dependent. **A:** Snapshots of MTs depletion visualised with SirTubulin upon activation of OptoDronc (Caspase 9) by light. Top row shows overlay of E-cad-tdTomato (green) and a clone expressing OptoDronc-GFP (magenta). Bottom row shows MTs using SirTubulin. White dotted line shows clone borders and the white arrow points an extruding cell. Scale bar is 5 μm. **B:** Schematic of the caspase activation cascade. OptoDronc is activated by 488nm light exposure through clustering and cleavage in trans, which then leads to effector caspase activation. p35 over-expresion inhibits effector caspase activity. **C:** Averaged and normalised medial apical sirTubulin signal (green) and perimeter (black) during cell extrusion induced by optoDronc. Light colour areas are s.e.m.. Grey dotted line represents the onset of extrusion. N= 1 pupae, n = 6 cells. **D:** Snapshots of UAS-optoDronc-GFP, UAS-p35 cells slowly contracting upon blue light exposure. Top row shows MTs accumulation visualised by SirTubulin. Middle row shows cells expressing UAS-OptoDronc-GFP and UAS-p35. Bottom row shows cell contour using E-cad-tdTomato. Scale bars are 5μm. **E,F:** Averaged and normalised medial apical sirTubulin signal (green), cell perimeter (black) (**E**) and averaged circularity (**F**, magenta) during the slow constriction of cell expressing UAS-OptoDronc-GFP and UAS-p35 upon blue light exposure. The pink curve shows the normal speed of extrusion upon optoDronc activation for comparison (see **C**). N= 2 pupae, n >99 cells. Light colour areas are s.e.m.. G: z-projection of a pupal notum showing cells stained for acetylated tubulin (pseudo-colour, right) and E-cad (green) inside and outside a clone depleted for the pro-apoptotic gene *hid* (UAS-hid-dsRNA, magenta). White lines: clone contour. White arrowheads point to some abnormal small cells in the clone showing an accumulation of acetylated tubulin (cell contour is shown with white dotted lines). Scale bar is 5μm.

### MT polymerisation/depolymerisation can increase/decrease cell apical area

The correlation between MT depletion and extrusion initiation (**Figure 3**), and the stabilisation of MTs observed upon caspase inhibition which prevents cell extrusion (**Figure 4**), suggest that MT disassembly may be permissive for apical area constriction and extrusion initiation. Accordingly, previous studies in the *Drosophila* pupal wing and embryo have shown a role of medio-apical MTs in cell apical area stabilisation^31,32^ We therefore tested whether MTs could also modulate cell apical area in the pupal notum. We injected colcemid (a MT depolymerising drug) in pupae and assessed the efficiency of MT depletion through the disappearance of EB1-GFP comets and the inhibition of mitosis progression (**Figure 5A**). UV exposure can locally inactivate colcemid^42^. Accordingly, local exposure to UV light (405nm diode) led to a rapid recovery of EB1 comets (**Figure 5A-A”, D, movie S15, bottom**). Strikingly, MT recovery was associated with a significant increase in cell apical area, which was not observed in the control regions, or upon UV exposure in mock-injected pupae (**Figure 5C-F, movie S15,** 4min after the onset of UV exposure). This suggested that local MTs polymerisation is sufficient to increase cell apical area on a timescale of minutes. lmportantly, the injection of colcemid did not lead to significant changes of MyoII levels during the first hours following injection, suggesting that these modulations are not driven by downstream effects on MyoII activation (**Figure S5 A-C**).

**Figure 5:**
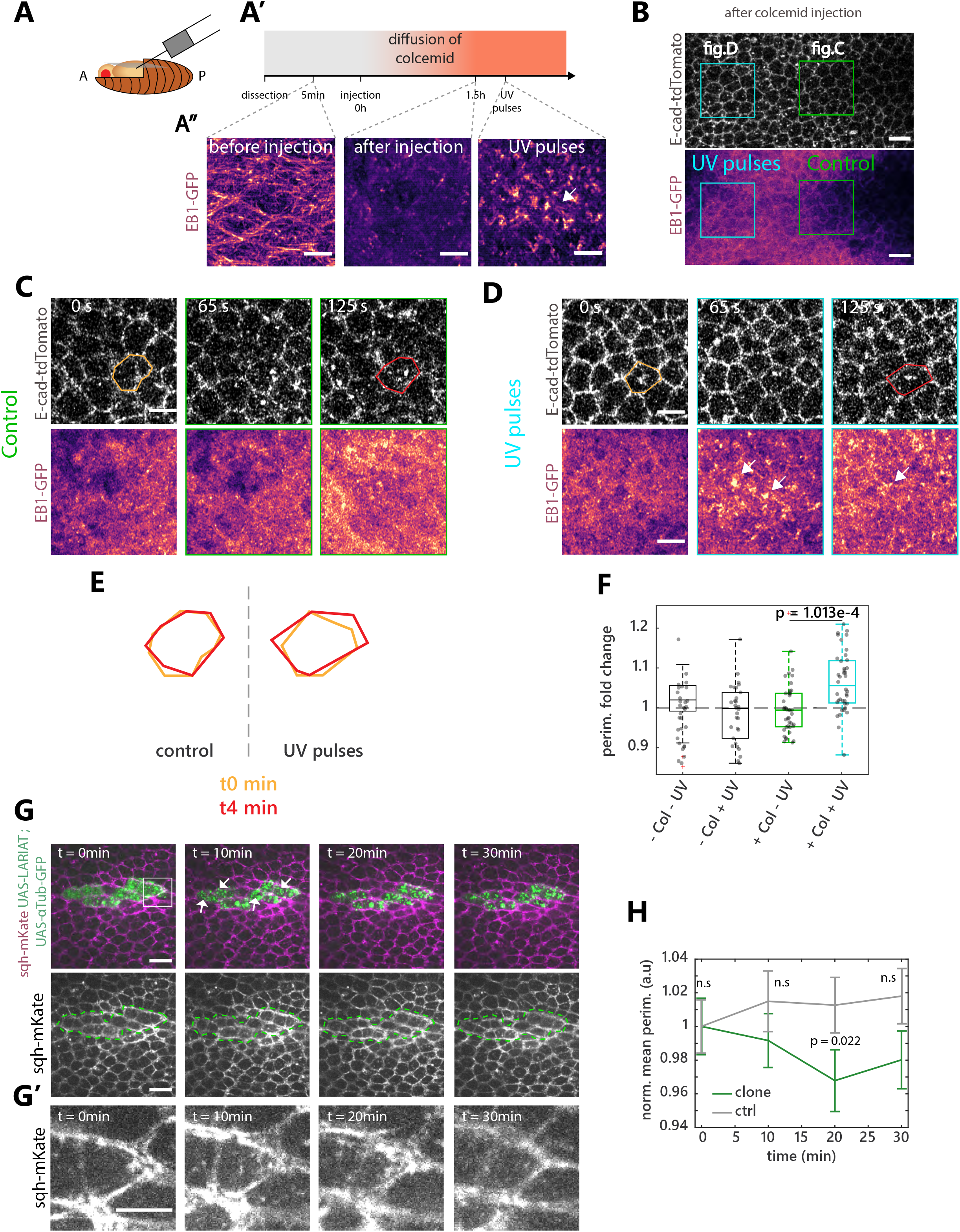
MTs polymerisation/depolymerisation can increase/decrease cell apical area. **A**: Schematic of the injection in the *Drosophila* pupae. **A’-A”**: Timeline of the injection protocol and colcemid inactivation experiment. Pupae were dissected and mounted between slide and coverslip to check for the organization of EB1-GFP before injection (see **A”** before injection time projection of EB1-GFP, pseudo-colour). Then pupae were unmounted and injected (see **Methods**) and remounted between slide and coverslip. We then waited for 1.5h for the drug to be active and checked again EB1 – GFP organisation (see **A”** after injection). We then pulsed UV in a selected region to inactivate colcemid locally (see **A”** UV pulses). Scale bars are 5μm **B:** Snapshot of the different experimental regions after colcemid injection, single plane. Top shows E-cad-tdTomato. Bottom shows EB1-GFP (pseudocolour, note the disappearance of MT structures). Green is the control region shown in **C.** Blue is the UV-exposed region shown in **D.** Scale bar, 10μm. **C:** Snapshots of a control region after colcemid injection. Top shows E-cad-tdTomato. Bottom shows EB1-GFP (pseudo-colour). Scale bars are 5μm. Orange and red contour, single representative cell at t0 and 4min. **D:** Snapshots of a region exposed to UV to inactivate colcemid 1.5h after colcemid injection. Top shows E-cad-tdTomato. Bottom shows EB1-GFP (pseudo-colour). White arrows point at foci of MT appearance. Scale bars are 5μm. Orange and red contour, single representative cell at t0 and 4min. **E:** Examples of cell size evolution during the experiment, orange t0, red 4min post UV exposition in the control region (left) or upon UV exposure (right), see **C, D.** Note the cell size increase from 0 to 4min upon UV exposure. **F:** Quantification of the cell perimeter fold change. Col stands for colcemid. Green box plot, control in condition with colcemid without UV pulses, blue box plot shows the condition with colcemid and UV. N = 4 pupae injected with colcemid and N = 3 control pupae, n > 30 cells per condition (one dot=one cell). **G-G’:** Snapshots of αTubulin-GFP clusterisation upon LARIAT activation and cell shape changes. **G.** Top, overlay of sqh-mKate (MRLC, purple, greyscale bottom) and a clone expressing UAS-LARIAT and UAS-αTubulin-GFP (green dotted line shows clone contours). Arrows point at the GFP clusters forming after exposure at 488nm. t0 is the start of the movie. The white box highlights the cell shown in **G’**. Scale bar is 10μm. **G’**: Snapshot of sqh-mKate of a cell upon LARIAT activation. Note cell constriction following 488nm exposure. Scale bar is 5μm. **H:** Normalised averaged perimeter upon LARIAT clusterisation of αTubulin-GFP (green) or in control cells (grey, outside the clone). n.s stands for not significant. N=3 pupae, n=41 cells for each time points.

We then tried to perform the reverse experiment (fast local depletion of MTs). Optogenetics can be used to trigger fast clustering and sequestration of proteins of interest. We used the LARIAT system (Light-Activated Reversible Inhibition by Assembled Trap) to trigger clustering of a GFP-tagged α-tubulin upon blue light exposure^43–45^. Since the GFP-tagged α-tubulin knock-in is not viable^46^, we instead used the overexpression of GFP-α-tubulin and LARIAT in clones to trigger a partial (endogenous Tubulin is still present) but significant depletion of α-Tubulin. Accordingly, blue light exposure led to rapid clustering of GFP α-Tubulin and a mild but reproducible reduction of cell apical area without clear modulation of MyoII (**Figure 5G-H, movie S16**). Altogether, we concluded that a fast and local increase (or decrease) of MTs is sufficient to expand (or respectively constrict) cell apical area independently of noticeable MyoII modulation.

### The disassembly of MTs by caspases is a rate-limiting step of extrusion

We then checked whether MT depletion was sufficient to trigger cell extrusion in the notum. Conditional induction of Spastin (a MT severing protein^47^) in clones was sufficient to increase the rate of extrusion, including in regions where no caspase activity is observed and where very few cells die in control conditions^18, 20, 21^ (**Figure 6A-B, movie S17**). To check whether MT depletion could indeed affect extrusion downstream of caspases, we assessed the impact of MT depletion on cells where caspase activation is inhibited. While the depletion of Hid by RNAi drastically reduces the rate of extrusion^20^, colcemid injection restored the rate of extrusion in *hid-RNAi* clones almost back to that of WT cells (**Figure 6C-D, movie S18**). lmportantly, while caspase activation almost systematically precedes cell extrusion in control conditions^18, 20, 21^, a large proportion of cells underwent extrusion in the absence of caspase activation (visualised with GC3Ai^21, 48^, a live sensor of effector caspase activity) in *hid RNAi* clones upon colcemid injection (**Figure 6E-F, movie S19**). This suggested that MTs depletion can bypass the requirement of caspase activation for cell extrusion. Accordingly, while the inhibition of effector caspases (using UAS-p35) combined with Caspase-9 activation (using optoDronc) drastically reduces the rate of extrusion and the cell constriction rate (**Figure 4D, E, movie S14 bottom**), the rate of constriction was significantly enhanced upon MT depletion by colcemid injection (**Figure 6G-H, movie S20**), albeit not back to WT speed. This confirmed that the accumulation of MTs we observed in optoDronc UAS-p35 clones (**Figure 4D, E**) is one of the factors slowing down cell constriction and cell extrusion. Altogether, this demonstrates that the disassembly of MTs by effector caspases is an essential rate-limiting steps of extrusion and one of the key initiators of cell constriction.

**Figure 6:**
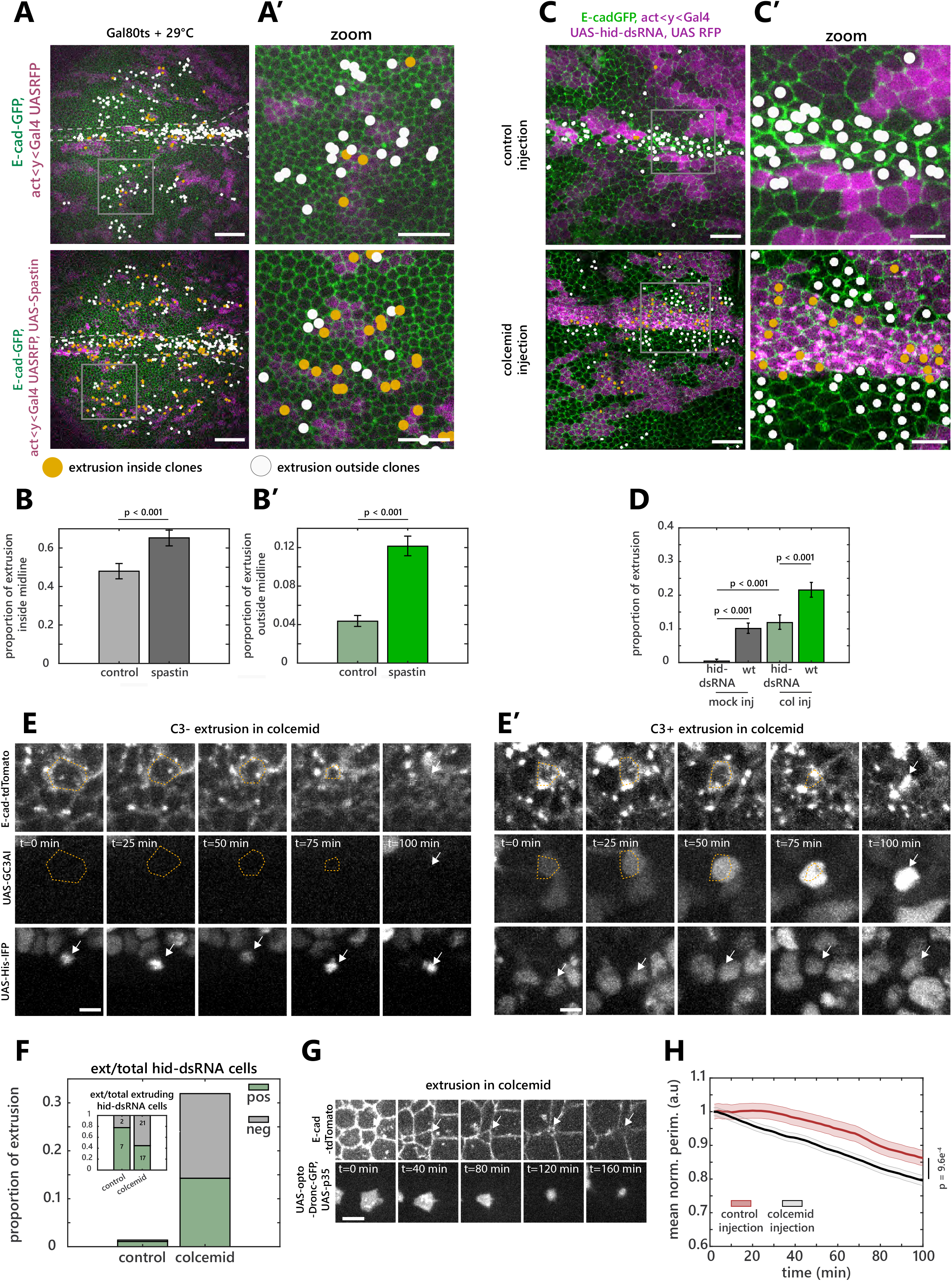
The disassembly of MTs by caspases is a rate limiting step of extrusion. **A-A’**. Live pupal nota expressing E-cad-GFP (green) and Gal4-expressing clones (magenta) driving the expression of RFP only (top) or the MT severing protein Spastin (bottom). Dots are marking the position of extrusion inside (orange) or outside clones (white) throughout the movie. White dotted lines show the midline. White rectangles shows the regions shown in A’. Scale bars are 50μm for **A** and 25 μm in **A’. B,B’:** Global probability of cell elimination in the presence of control clones or clones expressing Spastin, inside the midline (**B**) or outside the midline (**B’**). Note that the proportion was estimated by dividing over the total number of cells because of the difficulty to detect clonal cells at t0. The increase rate of cell elimination is therefore underestimated. Error bars are 95% confidence interval, p-values, Fisher exact test. n = 993 midline cells, n = 6997 cells outside midline from 3 independent pupae expressing spastin in clones. n = 630 midline cells, n = 5105 cells outside midline from 2 independent control pupae **C-C’:** Live pupal nota expressing E-cad-GFP (green) and clones depleted for *hid* (magenta, UAS-hid-dsRNA) upon injection of EtOH+H2O (top, control) or colcemid (bottom). Dots are marking the position of extrusions inside (orange) or outside clones (white) throughout the movie. Scale bars are 25μm for **C** and 10 μm in **C’**. Rectangles show the regions zoomed-in in **C’. D**: Proportion of cell eliminated inside hid-dsRNA clones and outside clones (wt) from nota mock injected (left) or upon colcemid injection (right). P-values from Fisher exact test. Error bars are 95% confidence interval. n = 1155 WT cells and n = 1150 cells from hid-dsRNA clones from 4 independent control pupae. n = 1388 WT cells and n = 926 cells expressing hid-dsRNA from 3 independent pupae injected with colcemid **E-E’**: Examples of cell extrusions upon colcemid injection in hid-dsRNA clones negative for effector caspase activity (**E**, GC3Ai, middle) or positive for effector caspase activity (**E’**). Histone-IFP (bottom) marks the hid-dsRNA cells. White arrows show the termination of extrusion and the nucleus of an extruding cell (bottom rows). Yellow dotted lines highlight the cell contour of the extruding cells. Scale bars are 5 μm. **F:** Proportion of extrusion positive or negative for caspase activation in control (mock injection) or upon colcemid injection in hid-dsRNA clones. N=2 pupae and n=9 extruding cells for mock, N=2 pupae and n=39 extruding cells for colcemid injected. The inset shows the proportion of each extrusion category in each condition with absolute numbers. **G:** Example of an extruding cell (white arrows) expressing OptoDronc-GFP and p35 upon colcemid injection (E-cad-tdTomato, top, optoDronc-GFP bottom). Scale bar = 5 μm. **H:** Averaged and normalised perimeter of cells expressing optoDronc and p35 upon blue light exposure in controls (red, see **Figure 5D-F**) or upon colcemid injection (black). Light colour areas shows s.e.m.. n>99 cells, N=3 mock injected pupae, N=2 colcemid injected pupae. Note that this quantification was performed on every cells, irrespective of the size of the optoDronc clone (which can affect the process of extrusion, see^21^).

## Discussion

Our quantitative characterisation of cell extrusion in the pupal notum led to the surprising observation that actomyosin dynamics are not sufficient to explain the early steps of cell extrusion. We found instead that the disassembly of an apical MTs network correlates with the initiation of cell apical constriction and cell extrusion, which is then followed by more typical actomyosin ring formation. This is, to our knowledge, one of the first descriptions of a role of MTs in the initiation of cell extrusion independently of MyoII. MTs have been previously shown to influence cell extrusion through their reorganisation in the neighbouring cells and the restriction of Rho activity toward the basal side^33^. Accordingly, we also observed an accumulation of MTs filaments in the neighbouring cells during the late phase of extrusion, which strikingly matches the timing of formation of a supracellular actomyosin ring. On the contrary, the novel function of MTs in the initiation of apical constriction that we described here is likely to be independent of Rho regulation. First, the initiation of cell deformation does not correlate with a change of actomyosin concentration/localisation/dynamics, nor a recruitment of Rho. Second, cells inhibited for caspases activation do not extrude, despite the significant accumulation of MyoII (upon depletion of Hid, **Figure S1H,H’** or using optoDronc combined with p35, **Figure 4D-G, Figure S4C**). Third, the evolution of apical cell shape during the early phase of extrusion does not match the evolution expected through an increase of line tension (unlike the extrusion of larval accessory cells, **Figure 2**). Altogether, this strongly argues for an initiation of extrusion which is independent of the formation of an actomyosin purse-string. As such, our study outlines a novel role of MTs in epithelial cell shape stabilisation which is independent of MyoII regulation. Recently, several works have described the central role of non-centrosomal MTs in epithelial morphogenesis, however this was mostly through their impact on MyoII activity^34–36^. Our study reinforces the notion that MTs may also stabilise cell apical area independently of MyoII regulation, as previously shown during morphogenesis of the *Drosophila* embryo^32^ or of the *Drosophila* pupal wing^31^. Interestingly, the accumulation of apical acetylated MTs promotes the capacity of cells to re-insert in an epithelial layer through radial intercalation in *Xenopus*^49, 50^ This nicely mirrors the function of MTs disassembly that we found in cell extrusion.

Through which mechanisms could MTs stabilise cell apical area? MTs are well known for their stabilising function of cell membrane protrusions during cell migration^51^ and can also modulate single cell major axis length^52^. Indeed, MTs embedded in the actomyosin network can bear significant compressive forces^53, 54^ and modulate cell compressibility^55^, or bear the compression driven by the constriction of cardiomyocytes^56^. Thus, apical MTs could directly resist the pre-existing cortical tension in the midline cells and their disassembly would be sufficient to trigger cell constriction. Alternatively, MTs may influence the contractile properties of the actomyosin cortex independently of Rho activity and MyoII phosphorylation, hence modulating cell deformation without apparent changes in actomyosin recruitment. Accordingly, MTs disassembly is sufficient to accelerate the kinetics of actomyosin constriction *in vitro*^57^. Finally, MTs may have a more indirect function either by modulating nuclei positioning^58, 59^, hence releasing space to facilitate apical constriction, or by directly modulating cytoplasmic viscosity^60^.

We found that MTs disassembly is driven directly or indirectly by effector caspases. The disappearance of apical MTs is not driven by a shift of the centrosome position^32^ (**Figure S3G**), nor a modulation of the localisation of the non-centrosomal MTs organisers Patronin and Shot^32, 35,36^ (**Figure S3E, F, movies S11,S12**). The disassembly seems instead to occur throughout the cell (**Figure S3H-I’**) and may be driven by a global modulation of core MTs components by caspases. Accordingly, α-tubulin and β-tubulin are both cleaved by caspases in S2 cells and these cleavages are conserved in humans^61^. This is in agreement with the depletion that we also observed with the tagged human α-tubulin (**Figure S3D**). However, we could not address the functional relevance of these cleavages since the mutant form of α-tubulin (mutation at the three cleavage sites) did not integrate properly in MT filaments either in S2 cells or in the notum (**Figure S7**). Since several core MT components are targets of caspases (including α-tub and β-tub)^61^, we believe that the inhibition of the caspasedependent disassembly of MTs will be hard to achieve. Of note, the redundancy of multiple caspase targets triggering MT depletion and the high conservation of several cleavage sites may reflect the physiological importance of this regulatory process.

We showed previously that caspase activation is required for cell extrusion in the pupal notum, including during cell death events induced by tissue compaction^18, 20, 21^. This suggested that cell extrusion is unlikely to occur spontaneously upon cell deformation and that permissive regulatory steps are required to allow cell expulsion. Our work suggests that the disassembly of MTs by caspases may be one key rate-limiting step. Accordingly, MTs depletion is sufficient to bypass the requirement of caspase activity for cell extrusion (**Figure 6**). Interestingly, several recently characterised mechanisms of MTs repair and stabilisation upon mechanical stress^38,39,62^ can reinforce the capacity of MTs to bear mechanical load^56^. Thus, stress generated by cell constriction and/or tissue compression is unlikely to be sufficient to trigger MT disassembly. This is in good agreement with the transient accumulation of MTs that we observed upon tissue stress release (**Figure S4A,B, movie S13**), and outlines the requirement for an active disassembly mechanism by caspases to trigger MT depletion and extrusion. It should be noted however that MT disassembly in caspase-inhibited cells does not completely rescue the speed and the rate of extrusion, thus other unidentified targets of caspases are likely to also participate in extrusion regulation.

Caspase activation in the pupal notum, and in other tissues, does not lead systematically to cell extrusion and cell death^18, 20, 21,63^ The mechanisms downstream of effector caspase activation governing cell survival or engagement in apoptosis remain poorly understood. Since the engagement of cells in extrusion in the WT notum systematically leads to cell death^17, 18^, and MT depletion is the earliest remodeling step associated with extrusion, the disassembly of MTs by caspases is likely to be one of the key decision steps leading to engagement in apoptosis in the pupal notum. Future work connecting cell mechanical state, quantitative caspase dynamics and MT remodeling may lead to important insights about the decision of a cell to die or survive.

## Acknowledgements

We thank members of RL lab for critical reading of the manuscript. We would like to thank Jakub Voznica for his observations on the abdomen during his internship. We are also grateful to Antoine Guichet, Thomas Lecuit, Magalie Suzanne, Yohanns Bellaïche, Xiaobo Wang, Renata Basto, the Bloomington Drosophila Stock Center, the Drosophila Genetic Resource Center and the Vienna Drosophila Resource Center, Flybase for sharing essential information, stocks and reagents. We also thank Benoît Aigouy for the Packing Analyser software and Jan Ellenberg group for MyPic autofocus macro. AV is supported by a PhD grant from the doctoral school “Complexité du Vivant” Sorbonne Université and from an extension grant of La Ligue contre le Cancer, work in RL lab is supported by the Institut Pasteur (G5 starting package), the ERC starting grant CoSpaDD (Competition for Space in Development and Disease, grant number 758457), the Cercle FSER and the CNRS (UMR 3738).

## Authors contribution

RL and AV discussed and designed the project and wrote the manuscript. AMV performed the vertex simulations. FL designed the tubulin mutant construct. AV performed all the other experiments and analysis. Every author has commented and edited the manuscript.

## Methods

### Ressource availability

#### Lead contact

Further information and requests for resources and reagents should be directed to and will be fulfilled by the lead contact, Romain Levayer.

#### Material availability

All the reagents generated in this study will be shared upon request to the lead contact without any restrictions.

#### Data and Code availability

All code generated in this study and the raw data corresponding to each figure panel (including images and local projection of movies) can be shared upon request and will be uploaded soon to a repository.

### Experimental model and subject details

#### Drosophila melanogaster husbandry

All the experiments were performed with *Drosophila melanogaster* fly lines with regular husbandry technics. The fly food used contains agar agar (7.6 g/l), saccharose (53 g/l) dry yeast (48 g/l), maize flour (38.4 g/l), propionic acid (3.8 ml/l), Nipagin 10% (23.9 ml/l) all mixed in one liter of distilled water. Flies were raised at 25°C in plastic vials with a 12h/12h dark light cycle at 60% of moisture unless specified in the legends and in the table below (alternatively raised at 18°C or 29°C). Females and males were used without distinction for all the experiments. We did not determine the health/immune status of pupae, adults, embryos and larvae, they were not involved in previous procedures, and they were all drug and test naïve.

#### Drosophila melanogaster strains

The strains used in this study and their origin are listed in the table below.

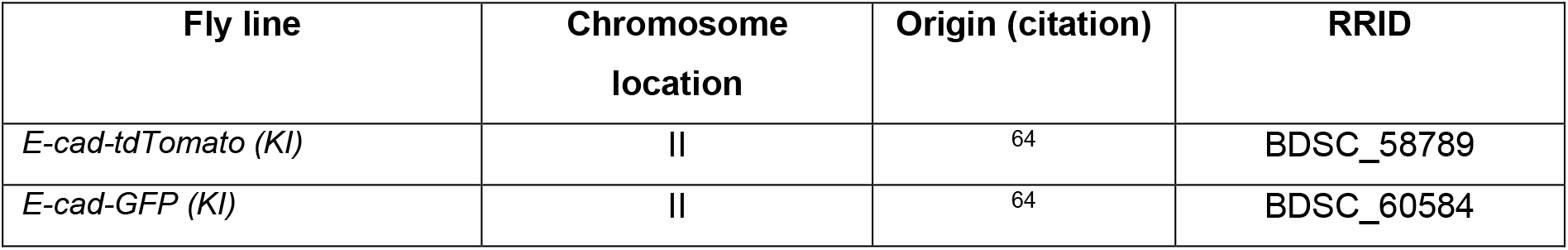
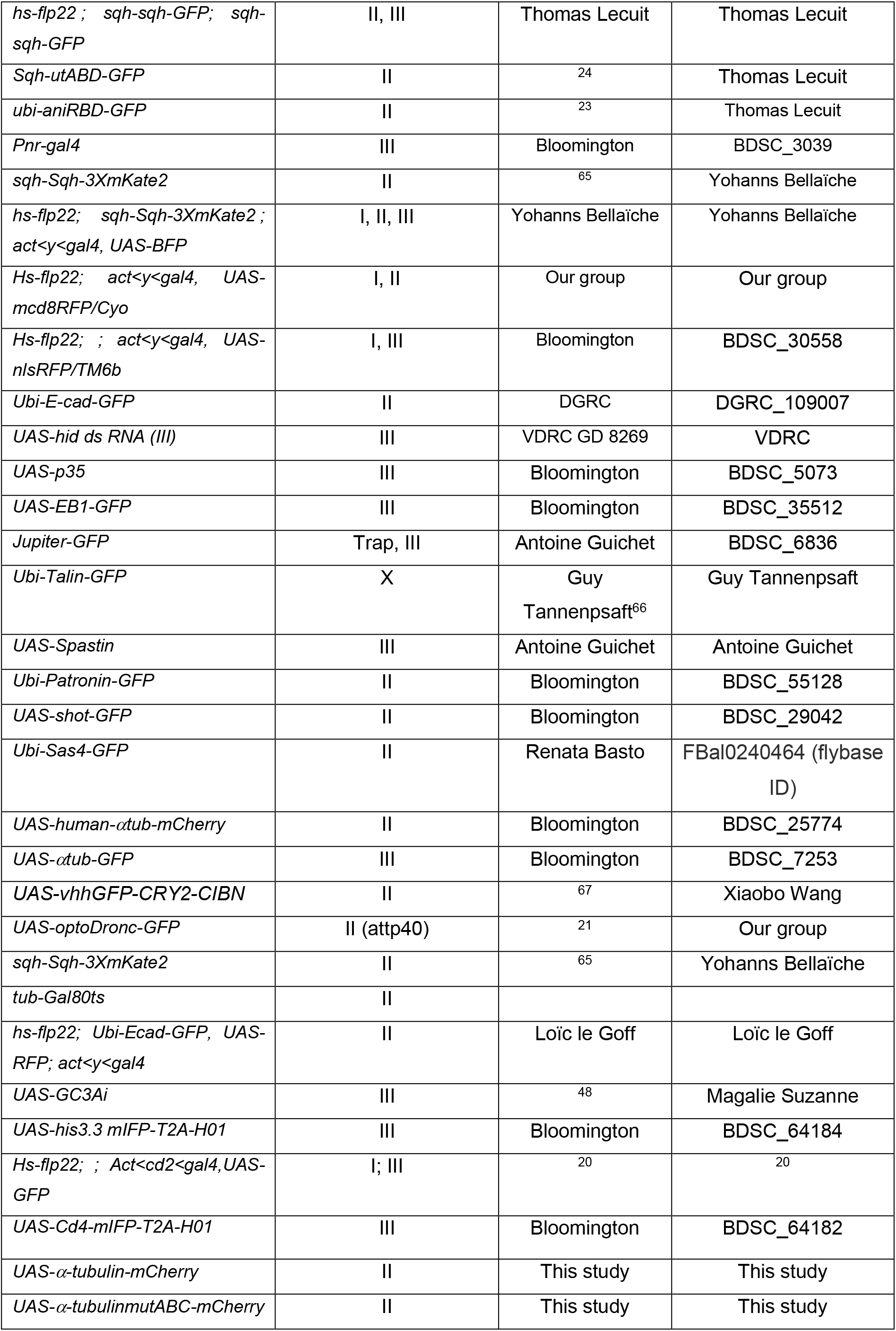

The exact genotype used for each experiment is listed in the next table. ACI: time After Clone lnduction, APF: After Pupal Formation, hs: heat shock at 37°C.

The exact genotype used for each experiment is listed in the next table. ACI: time After Clone Induction, APF: After Pupal Formation, hs : heat shock at 37°C.
| Figure | Genotype | Clone induction | Dev. time |
| --- | --- | --- | --- |
| 1 B | E-cad-GFP(KI); pnrGal4 | - | 16h APF |
| 1 C, D | hs-flp22; ubi-E-cad-GFP UAS-RFP/+;<br>act<y<Gal4/UAS-hid-dsRNA (quantification outside clones) | - | 16h APF |
| 1 E-F, I-J' ;S1 A-F | hs-flp22; sqh-Sqh::GFP; sqh-Sqh::GFP | - | 20h APF |
| 1 G-H, S1 G-H | hs-flp; act<y<gal4, UAS-mcd8RFP/Cyo ; sqh-GFP::utABD/TM6b | - | 16h APF |
| S1 I-J' | w; E-cad-tdTomato (KI)/+; +/ubi-mE ::AnilRBDGFP | - | 16h APF |
| S1 H-H' | yw, hs-flp22 ; sqh-sqh::GFP/Cyo ; act<cd2<gal4 , UAS-nlsRFP/UAS-hid-dsRNA | 48h ACI, 12min hs | 20h APF |
| 2 I | w; E-cad-GFP(KI) | - |  |
| 2 J | w; E-cad-tdTomato /+; pnr-Gal4/UAS-EB1-GFP | - |  |
| S2 A top | hs-flp22; sqh-Sqh::GFP; sqh-Sqh::GFP | - | 20h APF |
| S2 A bottom | w; E-cad-GFP(KI) | - | 20h APF |
| S2 B-C | w ; UAS-optoDroncCRY2-GFP/ E-cad-tdTomato; MKRS/act<y<Gal4 UAS-GFP | 48h ACI, 12min hs | 16h APF |
| S2 D | ubi-Talin::GFP/FM7 | - | 16h APF |
| 3 A, B' | w; E-cad-tdTomato | - | 16h APF |
| F3 C-G' | w; E-cad-tdTomato /cyo; Jupiter-GFP/TM6B | - | 16h APF |
| S3 A-C | w; E-cad-tdTomato(KI) /+; pnr-Gal4/UAS-EB1-GFP | - | 16h APF |
| S3 D | w; E-cad-GFP(KI) /+; pnr-Gal4/UAS-alpha-Tub-mCherry-human | - | 16h APF |
| S3 E | hs-flp22; E-cad-tdTomato (KI) /UAS-shot-GFP ; pnr-Gal4/+ | - | 16h APF |
| S3 F | w;ubi-p63-Patronin.A.GFP/Cyo | - | 16h APF |
| S3 G | w; E-cad-tdTomato(KI)/ubi-Sas4-GFP | - | 16h APF |
| S3 H-I' S4 A-B | w ; E-cad-GFP(KI); /UAS-α-tubulin-mCherry ; pnr-Gal4/+ | - | 16h APF |
| 4 A | w ; UAS-optoDroncCRY2-GFP/ E-cad-tdTomato(KI); MKRS/act<y<Gal4 UAS-GFP | 48h ACI, 12min hs | 16h APF |
| 4 D-F | w ; UAS-optoDroncCRY2-GFP/ E-cad-tdTomato; UAS-p35/act<y<Gal4 UAS-GFP | 48h ACI, 12min hs | 16h APF |
| 4 G | yw, hs-flp22 ; E-cad-tdTomato(KI)/Cyo ; act<cd2<gal4 , UAS-GFP/UAS-hid-dsRNA | 48h ACI, 12min hs | 20h APF |
| S4 A,B | w ; E-cad-GFP(KI); /UAS-α-tubulin-mCherry ; pnr-Gal4/+ | - | 30h APF |
| S4 C | w ; sqh-sqh-mKatex3/ UAS-optoDroncCRY2-GFP ; act<y<Gal4, UAS-BFP/UAS-p35 | 48h ACI, 12min hs | 30h APF |
| 5 A-F | w; E-cad-tdTomato(KI) /+; pnr-Gal4/UAS-EB1-GFP | - | 16h APF |
| 5 G-H | Hs-flp22; sqh-sqh-mKatex3/UAS-vhhGFP-CRY2-CIBN; act<y<Gal4, UAS-BFP / UAS-αTubulin-GFP | 48h ACI, 12min hs | 16h APF |
| S5 A-C | w ; sqh-sqh-mKatex3/Cyo ; pnr-gal4/TM6b | - | 16h APF |
| 6 A-B' | hs-flp22; tub-Gal80ts/ ubi-E-cad-GFP, UAS-RFP ; UAS-Spstin/act<y<Gal4 | 96h ACI 18°C, 12min hs | 16h APF |
| 6 A-B' | hs-flp22; tub-Gal80ts/ ubi-E-cad-GFP, UAS-RFP ; MKRS/act<y<Gal4 | 96h ACI 18°C, 12min hs | 16h APF |
| 6 C-C', D | hs-flp22; ubi-E-cad-GFP, UAS-RFP/+; act<y<Gal4/UAS-hid-dsRNA | 48h ACI, 12min hs | 16h APF |
| 6 E-F | hs-flp22 ; E-cad-tdTomato(KI) /UAS-GC3AI ;<br>act<y<Gal4, UAS-His3-mIFP/UAS-hid-dsRNA | 48h AIC,<br>12min hs | 16h APF |
| 6 G-H | hs-flp22 ; E-cad-tdTomato(KI)/ UAS-optoDroncCRY2-GFP ; act< y<Gal4, UAS-GFP/UAS-p35 | 48h AIC,<br>12min hs | 16h APF |
| S7 C-D | w ; E-cadGFP(KI)//UAS-α-tubulin-mCherry-ABC ;<br>pnr-Gal4/+ | - | 16h APF |
| S7 C-D | w ; E-cadGFP(KI)//UAS-α-tubulin-mCherry; pnr-<br>Gal4/+ | - | 16h APF |

#### Generation of α-tub84B-mCherry WT and non-cleavable mutant

The inserts mCherry-alphaTub84b and mCherry-alphaTubD34A-D48A-D200A (mutation of the three caspase cleavage sites) were generated by PCR from pAc-mCh-Tub (addgene 24288) with oligos containing mutations or not. The triple mutant at the three sites (mCherry-alphaTubD34A-D48A-D200A) was generated by using the following primers combination (see table below for primer sequence): F1+R1, F2+R2, F3+R3. The WT form was generated using the F1+R3 primers. The PCR products were then inserted in the pJFRC4-3XUAS-IVS-mCD8::GFP (Addgene 28243) linearized by NotI, XbaI digestion, using NEBuilder HiFi DNA Assembly Method. The construct was checked by sequencing and inserted at the attp site attp40A after injection by Bestgene. The primers used for the construct are listed below (inserted mutations sites are shown in red).

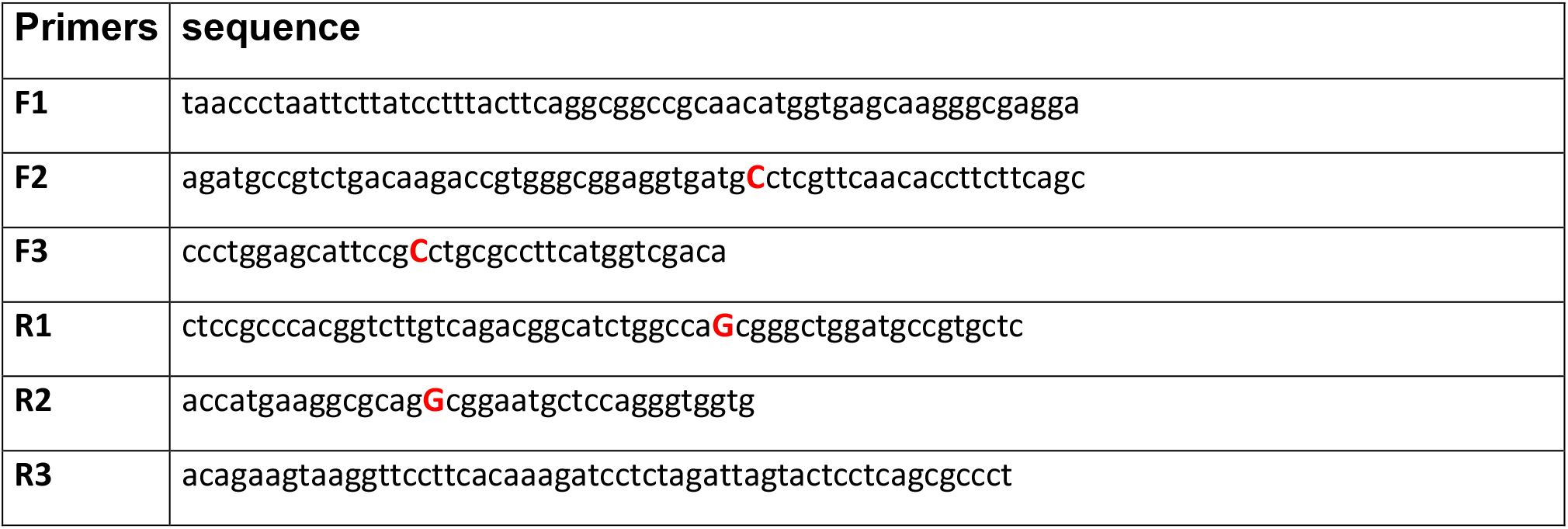

### S2 cell culture

S2R+ cells were cultured in Schneider’s *Drosophila* Medium with 10% fetal bovine serum, penicillin and streptomycin. S2R+ cell were transfected with FUGENE HD (Promega, ref: E2311). 24h after transfection, S2R+ cells were plated on glass-bottom dishes coated with concanavalin A (con A). Cells were imaged in a spinning disk confocal 30–60 min after cell spreading on the dishes.

### Vertex modeling of cell extrusion

To model the early steps of extrusion, we used a computational vertex model based on the existing computational framework for the study of developmental processes in the epithelial tissues of *Drosophila*^68, 29^. The model was implemented in gfortran, using openGL to visualize the outputs.

In the vertex model, only the apical sides of the cells are considered. Cells are represented as 2D polygons, made of vertices connected by edges. The vertices can move over time as a result of intra- and inter-cellular mechanical forces. The movement of the vertices is implemented by comparing the mechanical energy of a vertex in its current position (x, y) with the energy of a randomly chosen point nearby (x+δd, y+δd) with δd €[0,0.005]. When the energy in the new position is smaller, then the movement is accepted as the new vertex location. When the energy is bigger, the movement is accepted with probability P_accept_ (= 0.05) in order to introduce stochastic fluctuations.

The energy (E) of a vertex i is given by

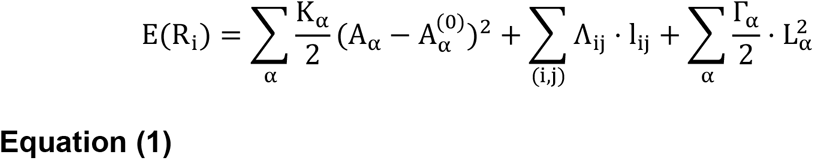

where R_i_ = (x_i_, y_i_) is the position of the vertex i. The first and the third summations are over all the cells α in which the vertex i is present, and the second summation is over all the cell edges {i, j} in which the vertex i is present. A_α_ is the apical area of the cell α and K is the area elasticity modulus, which is assumed to be equal for all the cells in our simulations. A_α_^(0)^ is the resting area of the cell α. The distance and the line tension between the pairs of vertices {i, j} are denoted l_ij_ and Λ_ij_, respectively. The third term includes the perimeter of the cell α (L_α_) and the perimeter contractility coefficient (Γ_α_). By choosing 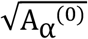 as a unit of length and (KA_α_^(0)^)^2^ as a unit of energy (as in^68^), dividing both sides of Eq. 1 by (KA_α_^(0)^)^2^ results in the following dimensionless equation:

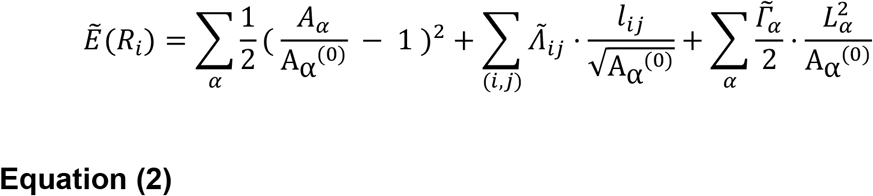

Where 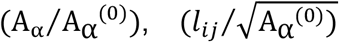 and 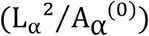 are, respectively, dimensionless area, bond length and perimeter. This model is characterized by dimensionless line tension 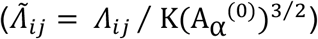 and dimensionless perimeter contractility 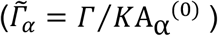 that were set respectively to 0.06 and 0.02 as in^69^.

Rearrangements of the topology of the vertices (T1 transitions) were allowed when two vertices i, j were located less than a minimum distance d_min_ (= 0.2) apart, and a movement of one of the vertices was energetically favorable such that the distance between the vertices decreases.

To avoid buckling at the boundary of the tissue, we assumed a greater stiffness of the cells edges located at the external boundary of the tissue, and set that the line tension for the external edges was higher 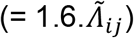 than that of the internal edges.

Simulations started from a tissue of 1141 cells, among which 10 cells scattered in the tissue (far from the edge to avoid boundary effects) were tracked for circularity (C = 4.π (area/perimeter^2^)) and perimeter (P) changes during the simulation run. In the initial topology, tracked cells have an average P of 7.58 ± 0.05 (s.e.m.) and an average C of 0.81 ± 0.088 (**Figure 2A**). Simulations were run for 6.5 million iterations, one iteration consisting in moving a randomly chosen vertex, updating its energy, and deciding to accept the movement or not. For clarity, simulation run was divided in 50 simulation time steps (sts), with 1 sts = 130.000 vertex iterations.

At t = 20 sts, three different conditions were examined to test for the effect of the mode of extrusion on cells circularity during the early stages of extrusion. In the first condition, the 10 tracked cells had parameters values identical to all the other cells of the tissue (control). In the second condition, the 10 tracked cells were forced to initiate extrusion by increasing at each iteration their contractility parameter 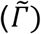 with a fixed rate 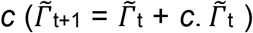 thus simulating a purse-string driven extrusion. Five different values were examined for *c*, with *c* = {0.0; 1.0; 2.5; 5.0; 7.5} x 10^-7^. In the third condition, the 10 tracked cells were forced to initiate extrusion by decreasing after each iteration their resting area (A_α_^(0)^) with a fixed rate *r* (A_α_^(0)^t+1 = A_α_^(0)^t – *r*. A_α_^(0)^t). Simulations were run for five *r* values (*r* = {0.0; 0.5; 1.0; 2.5; 3.5} x 10^-4^).

### Optogenetic control

#### Induction of cell death using optoDronc

For induction of optoDronc in clones in the pupal notum, *hs-flp; E-cad-tdTomato(KI); act<cd2<G4, UAS-GFP* females were crossed with homozygous *UAS-optoDronc* or *UAS-optoDronc; UAS-p35.* Clones were induced through a 12 minutes heat shock in a 37°C waterbath. Tubes were then maintained in the dark at 25°C. White pupae were collected 48 to 72 hours after clone induction and aged for 16h at 25°C in the dark. Collection of pupae and dissection were performed on a binocular with LED covered with home-made red filter (Lee colour filter set, primary red) after checking that blue light was effectively cut (using a spectrometer). Pupae were then imaged on a spinning disc confocal (Gataca system). The full tissue was exposed to blue light using the diode 488 of the spinning disc system (12% AOTF, 200ms exposure per plane, 1 stack/min). Extrusion profiles were obtained by segmenting extruding cells in the optoDronc clones with E-cad-tdTomato signal in the notum using Tissue analyzer ^70^. Curves were aligned on the termination of extrusion (no more apical area visible) and normalised with the averaged area on the first five points. The same procedure was used upon control injection or injection of colcemid (see below) in optoDronc UAS-p35 background. Note that in this condition, all cells of the clones were segmented irrespective of the size of the clone, which can affect by itself the speed of extrusion^21^.

#### LARIAT mediated depletion of αTubulin-GFP

UAS-vhhGFP-CRY2-CIBN (hereafter LARIAT) was expressed in gal4-expressing clones. CRY2 dimerises with CIBN in a light dependent manner. The association with the anti-GFP nanobody (vhhGFP) allows to trap the αtubulin-GFP in these large clusters. Clones were induced through a 12 minutes heat shock in a 37°C waterbath. Tubes were then maintained in the dark at 25°C. White pupae were collected 48 to 72 hours after clone induction and aged for 16h at 25°C in the dark. Pupae were then dissected an imaged using the same method than described for the optoDronc condition. Quantification were made by segmenting manually cells from clones and in control population at 4 time points (t = 0, 10, 15, 20 min).

### Live imaging and movie preparation

Notum live imaging was performed as followed: the pupae were collected at the white stage (0 hour after pupal formation), aged at 25° or 29° (glued on double sided tape on a slide and surrounded by two home-made steel spacers (thickness: 0.64 mm, width 20×20mm). The pupal case was opened up to the abdomen using forceps and mounted with a 20×40mm #1.5 coverslip where we buttered halocarbon oil 10S. The coverslip was then attached to spacers and the slide with two pieces of tape. Pupae were collected 48 or 72h after clone induction and dissected usually at 16 to 18 hours APF (after pupal formation). The time of imaging for each experiment is provided in the table above. Pupae were dissected and imaged on a confocal spinning disc microscope (Gataca systems) with a 40X oil objective (Nikon plan fluor, N.A. 1.30) or 100X oil objective (Nikon plan fluor A N.A. 1.30) or a LSM880 equipped with a fast Airyscan using an oil 40X objective (N.A. 1.3) or 63X objective (N.A. 1.4), Z-stacks (0.5 or 1 μm/slice), every 5min or 1min using autofocus at 25°C. The autofocus was performed using E-cad signal as a plane of reference (using a Zen Macro developed by Jan Ellenberg laboratory, MyPic) or a custom made Metamorph journal on the spinning disc. Movies were performed in the nota close to the scutellum region containing the midline and the aDC and pDC macrochaetae. Movies shown are adaptive local Z-projections. Briefly, E-cad plane was used as a reference to locate the plane of interest on sub windows (using the maximum in Z of average intensity or the maximum of the standard deviation) through the Fiji plugin LocalZprojector or corresponding MATLAB routine ^71^.

### Laser ablation

Photo-ablation experiments were performed using a pulsed UV-laser (355nm, Teem photonics, 20kHz, peak power 0.7kW) coupled to a Ilas-pulse module (Gataca-systems) attached to our spinning disk microscope. The module was first calibrated and then set between 30-40% laser power to avoid cavitation. Images were taken every 1min and ablation started after 1 time point. 400×400μm rectangle were converted to line of 10 thickness in metamorph. Repetitions were set between 5 and 10 for proper cut to be achieved. Cell perimeter was obtained through cell segmentation and the tubulin signal quantified in the total area of each cell (contour +3px).

### Image processing and inflection point detection

All images were processed using Matlab and FlJl (http://fiji.sc/wiki/index.php/Fiji). Movies for analysis were obtained after local Z projections of z-stacks using the Fiji LocalZprojector plugin^72^. As we were interested by apical signals we set ΔZ=1 so 3 planes of 0.5μm or 1μm were projected using maximum intensity projections. Then extruding cells were manually detected. When needed the signal was corrected for slight bleaching using CorrBleach macro from EMBL (https://www.embl.de/eamnet/html/bleach_correction.html). In order to measure signal intensities single cells were segmented using E-cad signal when it was possible (otherwise sqh or utABD signal). Depending on the data this was done directly using tissue analyzer after local z projections or after using epyseg (https://github.com/baigouy/EPySeg)^73^. Once segmented, ROI of the cell contour were extracted to Fiji and custom macro were used to measure the mean px intensities of medio-apical signal (−3px from the junction), total signal (+3px from the contour) or junctional signal (transformation to an area of 6px wide encompassing the junctions). Perimeter was measured using the real cell contour. Result were then analyzed in MATLAB.

For analysis single curve were aligned either by the end (i.e. the moment of the end of extrusion) or by the inflection point of the perimeter. Inflection points were automatically detected using a homemade MATLAB function. Briefly, it uses 2 moving linear fits after smoothing the perimeter using MATLAB moving average (taking into account 5 data point windows). The point with minimal error between the 2 fits and real data correspond to the point where the perimeter starts to constrict i.e. the inflection point. Average, standard deviation and s.e.m. were calculated after the alignment.

Radial averaged kymograph were obtained by tracking the centroid of every extruding cells and measuring the intensity along concentric circles of 3px at different distances from the cell center. The values were averaged for every tracked extruding cells. Ecad signal was used to detect cell contour and define MTs signal from the extruding cell and from the neighbours.

### Myosin peak detection and yield computation and cross correlation

We first computed the contraction rate as following 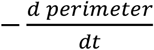 and the myosin rate of change as 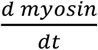. In order to assess how closely changes in myosin relate to constriction we computed the cross-correlation between myosin level or myosin rate of change and the constriction rate. The cross-correlation was calculated on Matlab with the *xcorr* function with the *‘coef’* option (normalized cross-correlation after subtracting the mean). All the curves (one per cell) were then aggregated and averaged.

Contraction peak and myosin peak were detected using the findpeaks function in Matlab by setting ‘MinPeakProminence’ to 7 in order to filter for noise. The yield was calculated for each contraction peak of each single curve as follow 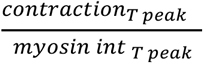. Perimeter curves were temporally aligned by their inflection point and yield data were sorted relative to the onset in 5min time windows. Then single curve were aggregated and averaged. In order to compute myosin pulse duration, amplitude and frequency we detected myosin pulses. We then returned the peak parameters: T_peak_ (Time at peak maximum), W_peak_ (width at half peak maximum) and A_peak_ (peak amplitude). Myosin pulse frequency was computed as follow: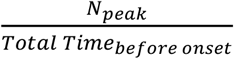 or 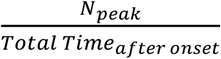.

### Drug injection in pupal notum

16h APF pupae were glued on double sided tape and the notum was dissected with red filter. Pupae were then injected using home-made needles (borosilicate glass capillary, outer diameter 1mm, internal diameter 0.5mm, 10cm long, Sutter instruments) pulled with a Sutter instrument P1000 pipette pulling apparatus. Colcemid (Sigma Demecolcine D7385 5mg) was diluted in ethanol to a stock concentration of 20mM and was then injected at 2.5mM in the thorax of the pupae using a Narishige IM400 injector using a constant pressure differential (continuous leakage).

### Colcemid inactivation by UV and effect of MT re-polymerisation on cell area

Pupae were dissected and mounted as described above. We first imaged EB1-GFP to assess its dynamics prior to colcemid injection. Then pupae were unmounted and we injected colcemid as described in this protocol. We then waited 1h30 for colcemid to diffuse in the notum. Next we re-imaged EB1-GFP and assessed colcemid effect through the loss of EB1-GFP comets and diffusion of the GFP pool as well as cell division arrest. (Cell division arrest was later used to assess colcemid effect whenever we could not image EB1-GFP signal).

We then image a single z-plane every second for 240s. We inactivated colcemid locally by pulsing 405nm diode (0.44% AOTF) in a restricted region of the imaging field and compared this to a control region. We assessed MT re-polymerisation by looking at the formation of EB1-GFP foci and comets. In order to measure the effect of repolymerisation on cell size we segmented the cell in the UV or control regions at t0 and t240s. As we are interested in the relative changes of cell perimeter between these two time points we computed a perimeter fold change for each cell as the following ratio 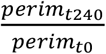 and compared the condition with colcemid to a control condition injected with H20 + ethanol.

### Proportion of cell elimination in clones

#### Spastin clones

Since we could not recover clone upon Spastin overexpression, we generated clones allowing conditional induction of Spastin with act<y<Gal4 UASRFP, UAS-Spastin with tub-Gal^80ts^. Gal^80ts^ binds Gal4 and represses Gal4-driven expression. Upon switching to 29°C Gal^80ts^ becomes inactive allowing Gal4-driven expression.

Due to the maturation time of RFP following the temperature shift to 29°C, it is difficult to track the position of the clones initially following temperature shift. For that reason we decided to measure the proportion of cell extrusion by looking at the global rate of cell extrusion at 29°C in the condition with control UAS-RFP of UAS-Spastin, UASRFP clones. Because of the high rates of cell extrusion in the midline we separated the quantification between the inside of the midline and outside. We did that by tracing manually the midline using the position of the most central Sensory Organ Precursors which define the midline. We then manually detected all the extrusions over 1000min and defined automatically if they belong to the midline or not. We then segmented the tissue at t0 to count the number of cells inside the midline or outside and then used these values to compute the proportion of extrusion as following: 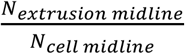 or 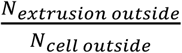.

#### Rescue of extrusion in hid-dsRNA following colcemid injection

For these experiments colcemid was injected as described above and this condition was compared with control injections (H20 + ethanol). We then manually detected all the extrusions at each time points and for each condition during the 500 first minutes. Clones were segmented at each time point using the UAS-RFP signal and we used this segmentation to automatically define if extrusions belong to UAS-hid-dsRNA 3 clones or not. We then segmented the tissue at t0min to count the number of cells in 3 the clones or outside the clones and used these values to compute the proportion of extrusion inside or outside the clone as following 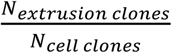 or 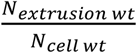.

#### Detection of Caspase signal in hid-dsRNA clones

Colcemid was injected as described previously in this protocol and this condition was compared with control injections (H20 + ethanol). We then manually detected all the extrusions at each time point and for each condition and manually defined if they are positive or negative for caspase activation using the UAS-GC3AI-GFP signal. We then 3 either computed the proportion of each ‘type’ of extrusion relative to the total number of cell in the clones or to the total number of extrusions in the clones.

### Statistics

Data were not analysed blindly. No specific method was used to predetermine the number of samples. The definition of n and the number of samples is given in each figure legend and in the table of the Experimental model section. Error bars are standard error of the mean (s.e.m.). p-values are calculated through t-test if the data passed normality test (Shapiro-Wilk test), or Mann-Whitney test/Rank sum test if the distribution was not normal, or Fisher exact test for comparison of proportion (see legends). Statistical tests were performed on Matlab.

---

**Figure S1:**
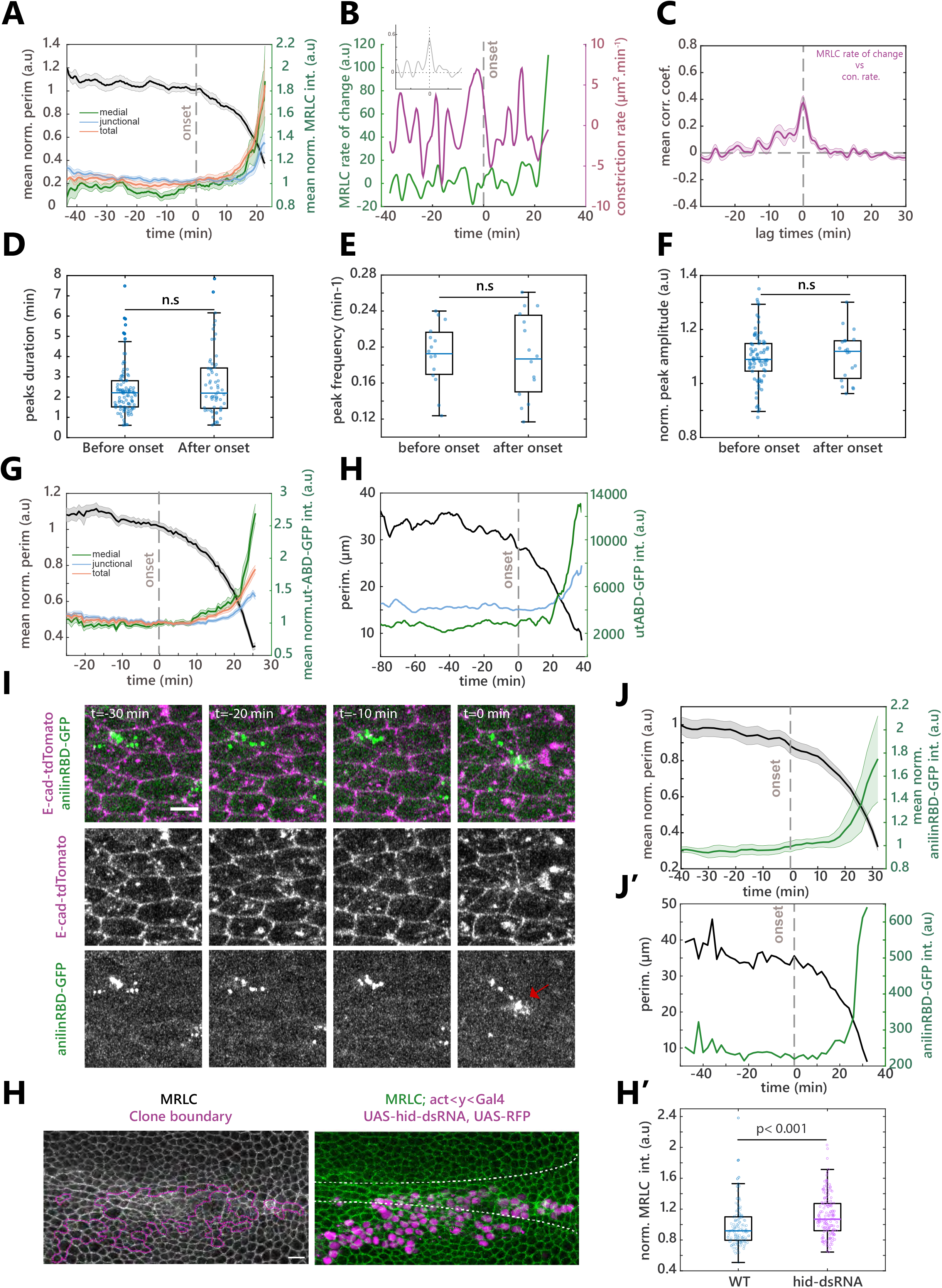
Myosin concentration and dynamics do not change at the onset of extrusion. **A:** Averaged and normalised intensity of MRLC (sqh-GFP) in different pools (medial in green, junctional in blue, total in orange) and perimeter (black) in midline extruding cells. Grey dotted line represents the onset of extrusion. Light colour areas are s.e.m.. n = 15 cells. **B:** Representative curve of MRLC (sqh-GFP) intensity rate of change (green) and perimeter contraction rate (purple) in a single extruding cell. Inset shows the normalized cross correlation of these two curves. t0 is the onset of extrusion (grey dotted line). **C:** Average normalised cross correlation of MRLC rate of change with perimeter contraction rate. n=15 cells. Light area is s.e.m.. **D,E,F:** Boxplots showing MRLC peaks duration (half peak width, **D**, n=106 & 61 pulses), peak frequency (**E**, n=15 cells) and normalised peak amplitude (**F,** n>70 and 20 peaks, before or up to 10 min after extrusion onset), before and after the onset of extrusion. n.s.: non-significant. **G:** Averaged normalised actin intensity (utABDGFP) in the medial (green), junctional (blue) and total pool (orange) and averaged normalized perimeter (black). Light colour areas are s.e.m.. Grey dotted line represents the onset of extrusion. n=37 cells. **H**: Representative curve of junctional (blue) and medial (green) actin (utABD-GFP) pools and perimeter (black) of a single extruding cell. **I:** Snapshots of Anillin Rho-Binding domain fused to GFP during cell extrusion (Rho sensor, green and bottom row) and E-cad-tdTomato (magenta and middle). Red arrow shows the late Rho accumulation matching the timing of the late actomyosin ring formation. Scale bar, 5μm. **J:** Averaged normalised AnilinRBD-GFP apical signal (green) and perimeter (black) during cell extrusion. Grey dotted line is the extrusion onset. Light colour areas are s.e.m.. N=2 pupae, n=31 cells. **J’**: Single representative curves of AnnilinRBD-GFP intensity (green) and perimeter (black) of a single extruding cell. **H:** Projection of MRLC-GFP (sqh-GFP, green and left) in a pupal notum with clones resistant for apoptosis (hid-dsRNA, magenta, borders shown with magenta lines). White doted lines show the midline. Scalebar is 10μm. **H’**: Boxplot of MRLC-GFP junctional intensity (one dot=one cell) outside or inside hid-dsRNA clones. N=2 pupae, n =779 WT cells and 155 clonal cells (note: only a subset of points are represented for the sake of visibility).

**Figure S2:**
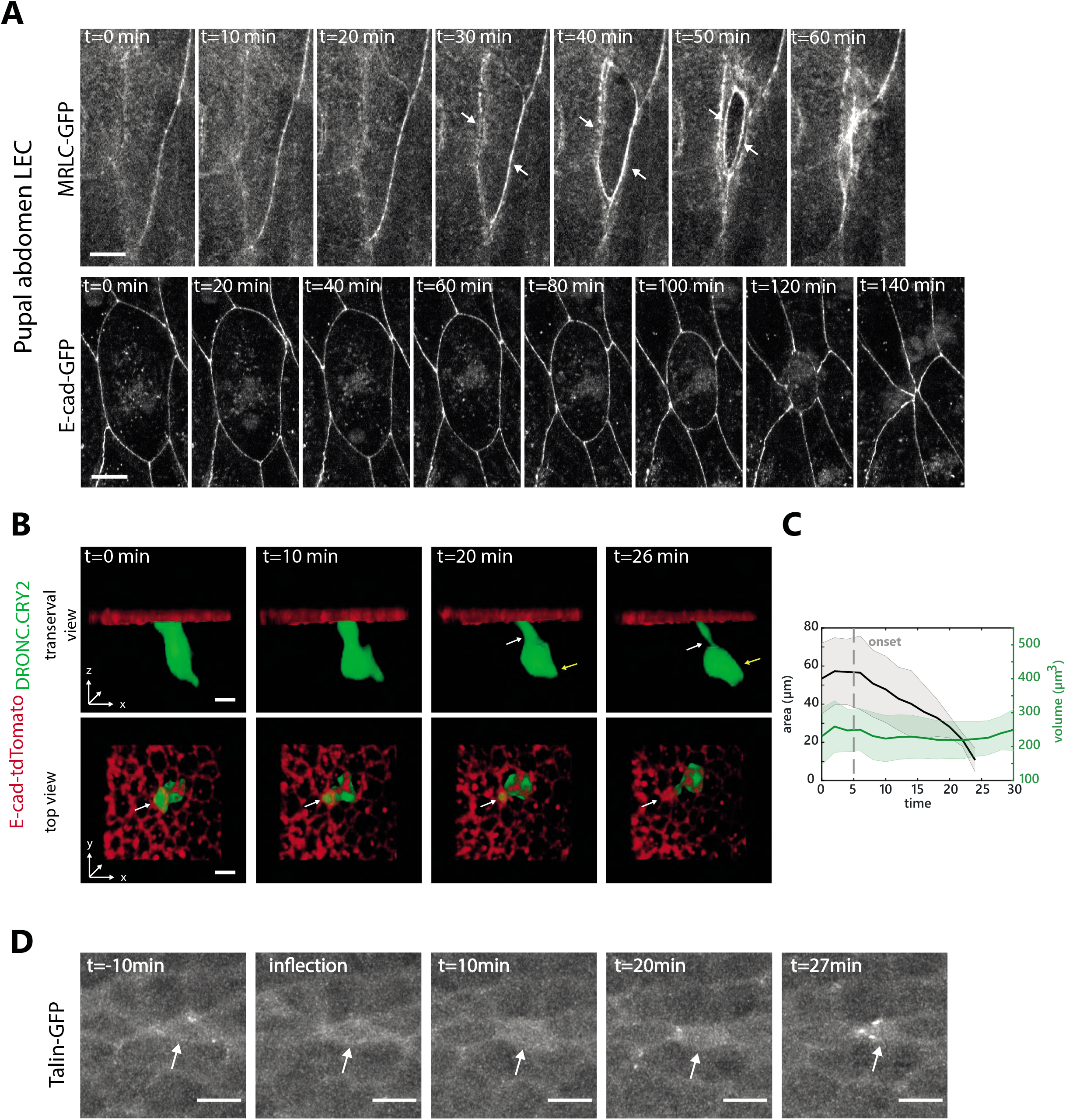
Cell extrusion in the notum is not initiated by an actomyosin ring, nor volume reduction or a modulation of ECM binding. **A:** Snapshots of extruding larval epidermal cells (LEC) from the pupal abdomen. Top shows MRLC (sqh-GFP). White arrows point at the supracellular and intracellular myosin ring. Bottom shows the evolution of cell perimeter visualised with E-cad-GFP which shows cell rounding. Scalebar is 10μm. **B:** Snapshots of a single extruding cell upon optoDronc activation with 3D rendering (using cytoplasmic GFP signal). Green is segmented cell volume, red is cell contour visualised with E-cad-tdTomato. Top row is a transversal view and bottom row a view from the apical side. White arrows point at apical reduction of volume while yellow arrow show basal increase of volume. Scale bars are 5μm. **C:** Averaged cell volume (green) and perimeter (black) during cell extrusion. Light colour areas are s.e.m.. n=6 cells. **D:** Snapshots of Talin-GFP during cell extrusion. t0 is the onset of extrusion (inflection). White arrows show an extruding cell. Scale bars are 5μm.

**Figure S3:**
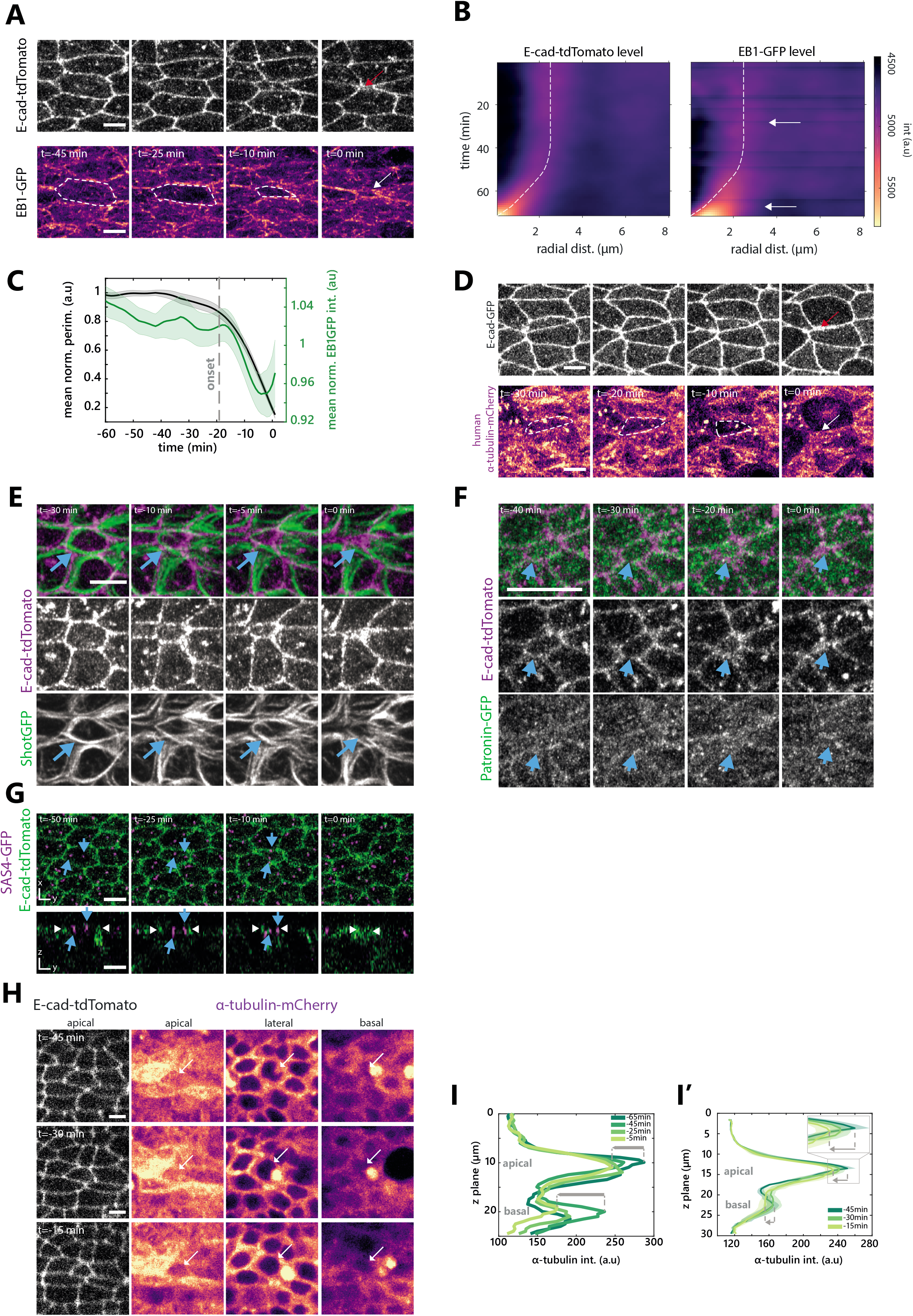
Visualisation of microtubule depletion using different markers. **A:** Snapshots of E-cad-tdTomato (top) and EB1-GFP (bottom, pseudo-colour) during cell extrusion in the midline. Red arrow points at the end of extrusion, white arrow shows late EB1 accumulation in the neigbours, the cell contour is shown with white dotted lines. Scale bar is 5μm. **B:** Radial averaged kymograph (see **Figure 3F-G’**) of E-cad-tdTomato (left) and EB1-GFP (right) in pseudo-colour, time is on the y-axis going downward, x-axis is radial distance from cell center. White dotted lines represent the average cell contour (localised by the local maximum of E-cad signal for each time point). Top white arrow shows the onset of EB1 depletion in the extruding cells, bottom arrow points at the late EB1-GFP accumulation in the neighbouring cells. n=60 cells **C:** Averaged normalised EB1-GFP signal (green) and perimeter (black). Grey dotted line represents the onset of extrusion. Light colour areas are s.e.m.. n=22 cells. **D:** Snapshots of E-cad-GFP (top) and human αTubulin-mCherry (bottom, pseudo-colour) during cell extrusion. Red and white arrows point at the end of extrusion. White doted lines show extruding cell contours. Representative of 15 cells. Scale bar is 5μm. **E:** Snapshots of Shot-GFP (green, bottom) and E-cad-tdTomato (magenta, middle) in an extruding cell. t0 is the termination of extrusion. Blue arrows show Shot-GFP filaments that persist over the course of cell extrusion. Representative of 20 cells. Scale bar is 5μm. **F:** Snapshots of Patronin-GFP (green, bottom) and E-cad-tdTomato (magenta, middle) in an extruding cell. t0 is the termination of cell extrusion. Blue arrow shows Patronin-GFP signal that persist over the course of cell extrusion. Representative of 20 cells. Scale bar is 10μm. **G:** Snapshot of centrosomes position during cell extrusion visualised by SAS4-GFP (magenta) with E-cad-tdTomato (green) in top and lateral view (bottom). t0 is the termination of cell extrusion. Blue arrows point at centrosome positions in the extruding cell (note the absence of movement along apico-basal axis). White arrows show the junctions of the extruding cell. Representative of 15 cells. Scale bar is 5μm. **H:** Snapshots of αTub-mCherry in different z-planes during cell extrusion (apical, leteral and basal, time from top to bottom, t0 is the termination of extrusion). Left panels show E-cad-tdTomato in the junction plane. White arrows point at the extruding cell in every plane. Scale bars are 5μm. **I-I’**: Quantification of apico-basal αtubulin-mCherry signal during extrusion (y-axis, z-plane, x-axis signal, different colour for each time point, t0 extrusion termination). **I:** Single representative quantification for the cells shown in **H**. Grey arrows show the reduction of signal over time apically and basally. I’: Averaged quantification of αtubulin-mCherry apico-basal signal during cell extrusion. Axis are the same than in I. The inset highlights apical depletion. Grey arrows show signal reduction in the apical and basal plane. n=11 cells

**Figure S4:**
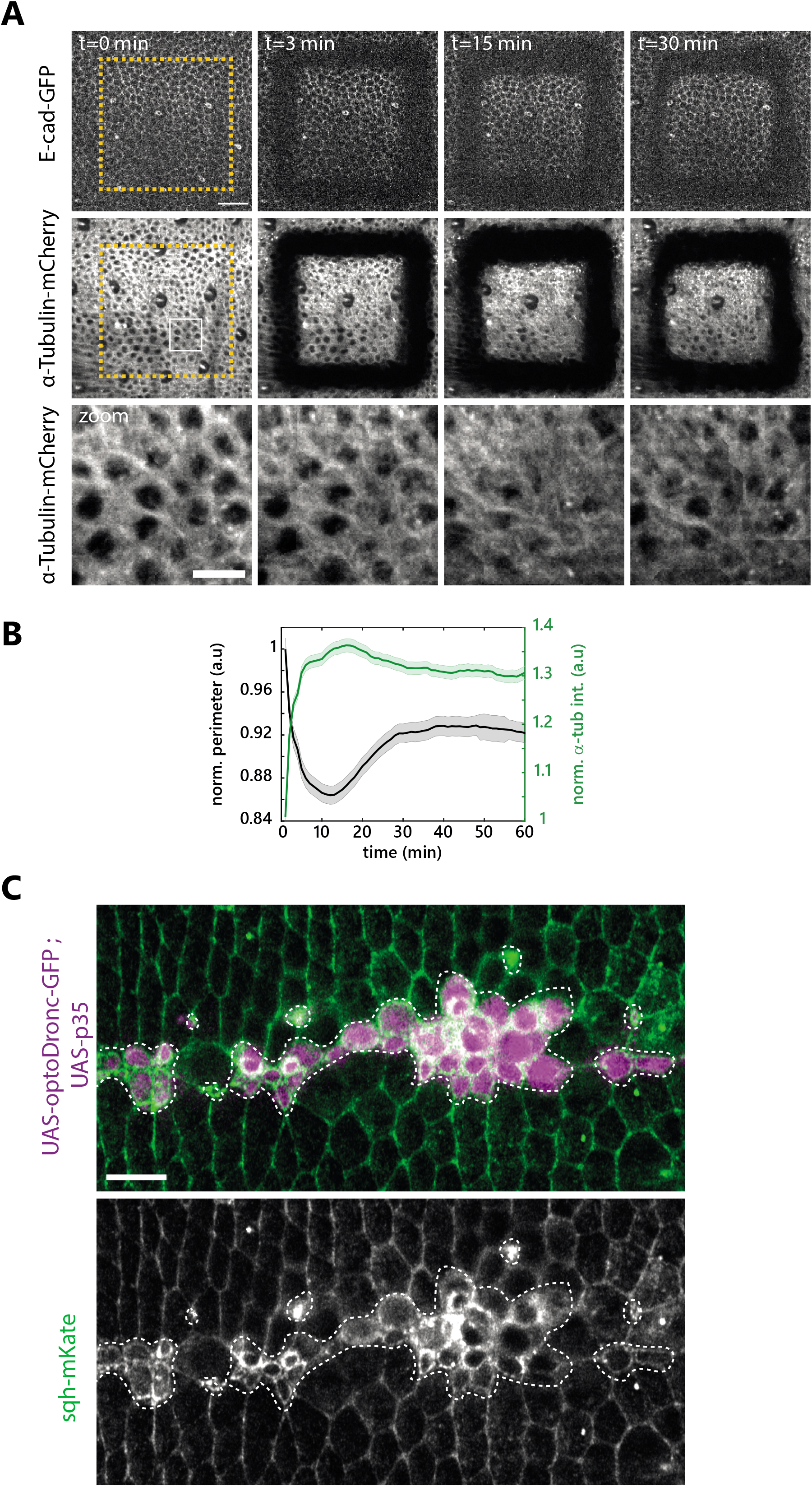
The depletion of microtubules is effector caspases dependent. **A:** Snapshots of a laser ablated pupal notum (orange dotted squares) showing α-Tubulin-mCherry (middle and bottom) and E-cad-GFP (top). t0 is the last time point before ablation. Scale bar is 25μm. The bottom row shows a zoom of α-Tubulin-mCherry from the white rectangle. Scale bar is 10μm. **B:** Averaged and normalised α-Tubulin-mCherry signal (green) and cell perimeter (black) in the relaxed square region following ablation. Note the transient increase of Tubulin signal during the transient reduction of cell perimeter. Light colour areas are s.e.m.. N=3 pupae, n=123 cells. **C:** Snapshot of MRLC accumulation (sqh-mKate, green and bottom) in clone expressing UAS-OptoDronc-GFP and UAS-p35 (magenta, contours shown with white dotted lines) after 3hours of blue light exposure. Representative of >10 clones. Scale bar is 10 μm.

**Figure S5:**
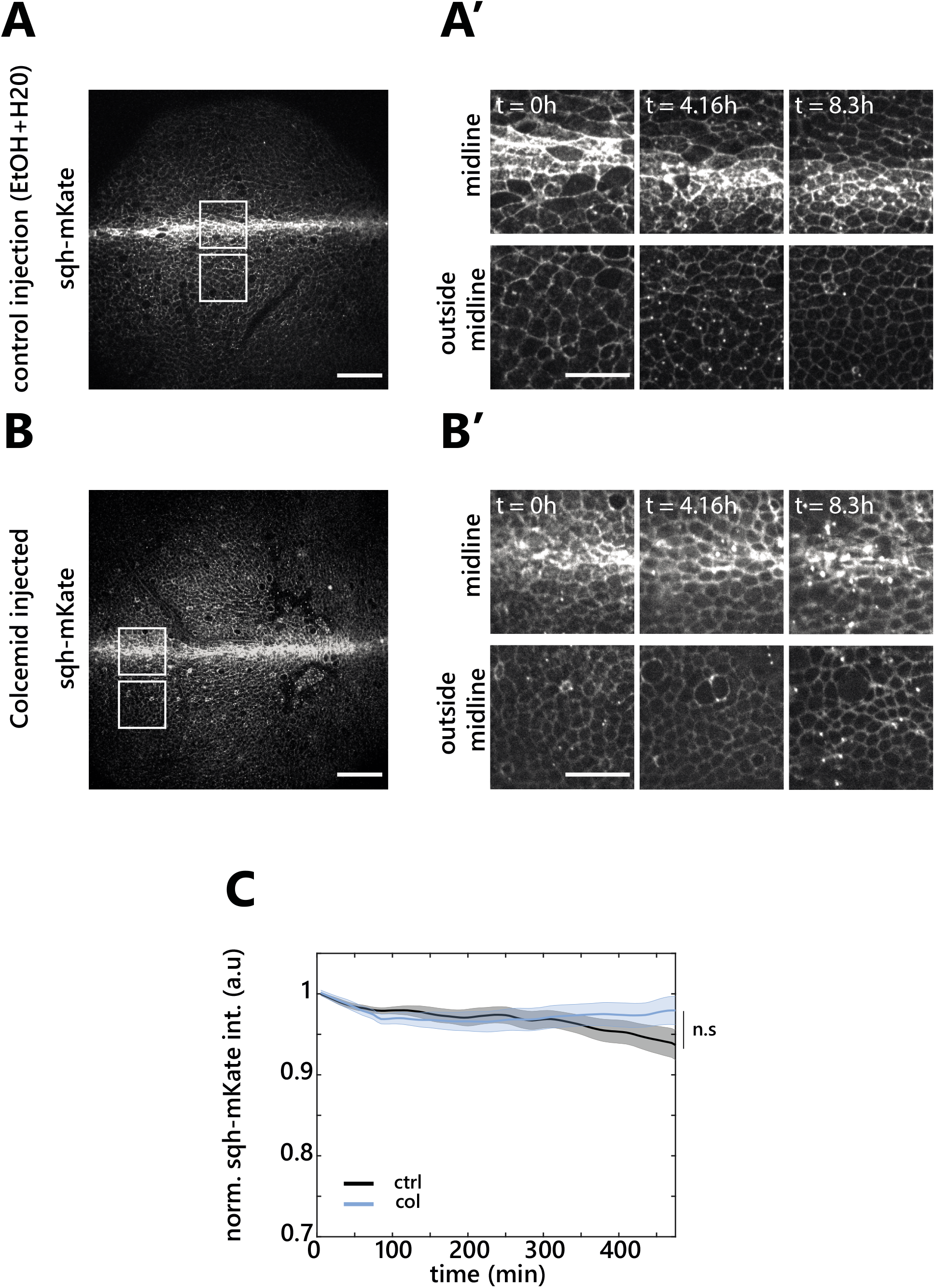
Evolution of MRLC levels upon colcemid injection. **A-B’.** Snapshots of local z-pojections of live pupal nota of Sqh-mKate (MRCL) levels following mock injection (EtOH+H20) (**A-A’**) or colcemid injection (**B-B’**) with the same acquisition parameters and contrasts. **A,B:** Full nota view right after injection, Scale bars are 50μm. White boxes highlight the midline (top) and out-of-midline (bottom) regions shown in **A’** and **B’. A’,B’:** Snapshots over time after injection. Scale bars are 25μm. **C:** Quantification of normalised and averaged Sqh-mKate signal from total fluorescence of 3 regions per pupae after injection in colcemid (blue) or control (black) injections. Light colour areas are s.e.m.. N=2 pupa for control and 3 pupa for colcemid.

**Figure S6:**
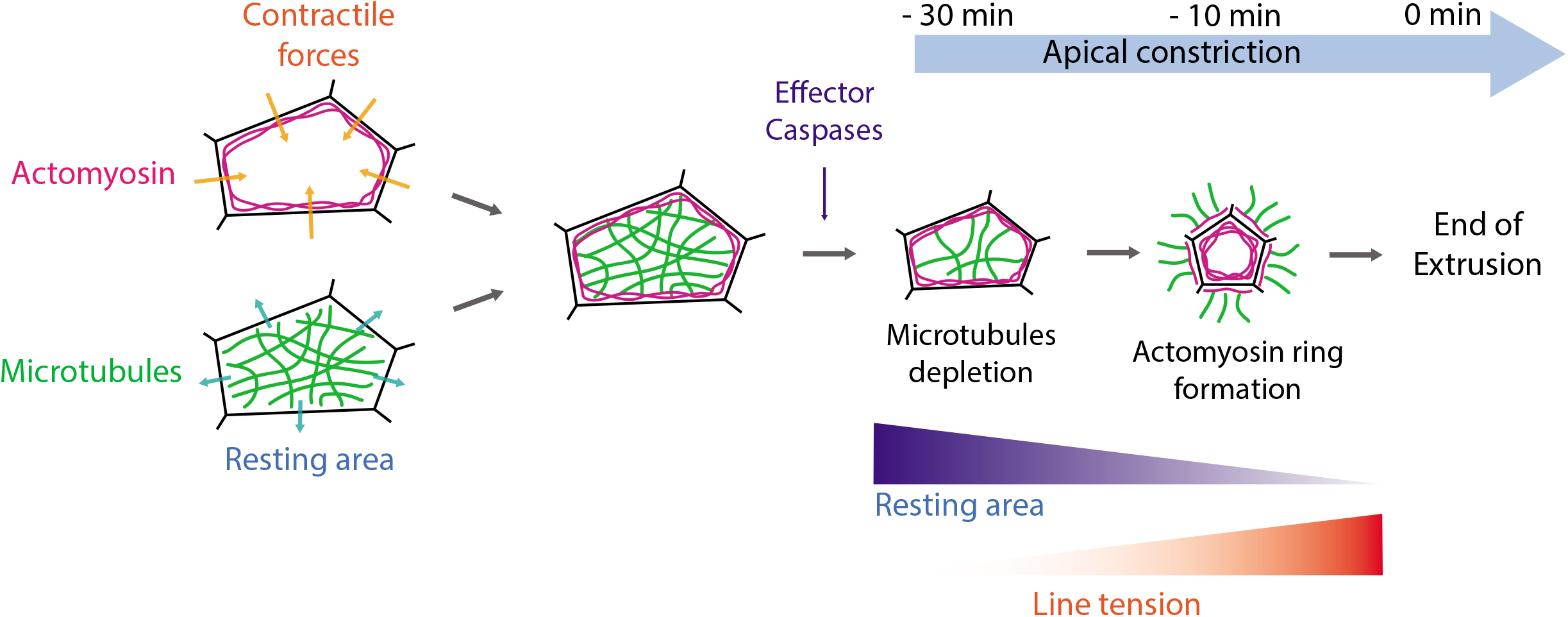
Working model. Schematic of the working model. Cell-cell junctions are in black. Actomyosin is in light red and microtubules (MTs) are represented as green lines. Contractile forces are regulated by the junctional actomyosin network. MTs may regulate directly or indirectly the resting area of the cell. At equilibrium the two components are balanced. Upon effector caspases activation, MTs are progressively depleted, hence reducing the resting area of the cell and promoting apical constriction without cell rounding. Later on, actomyosin accumulates in a contractile ring which terminates cell extrusion through an increase of line tension and correlating with an accumulation of MTs in the neighbours.

**Figure S7:**
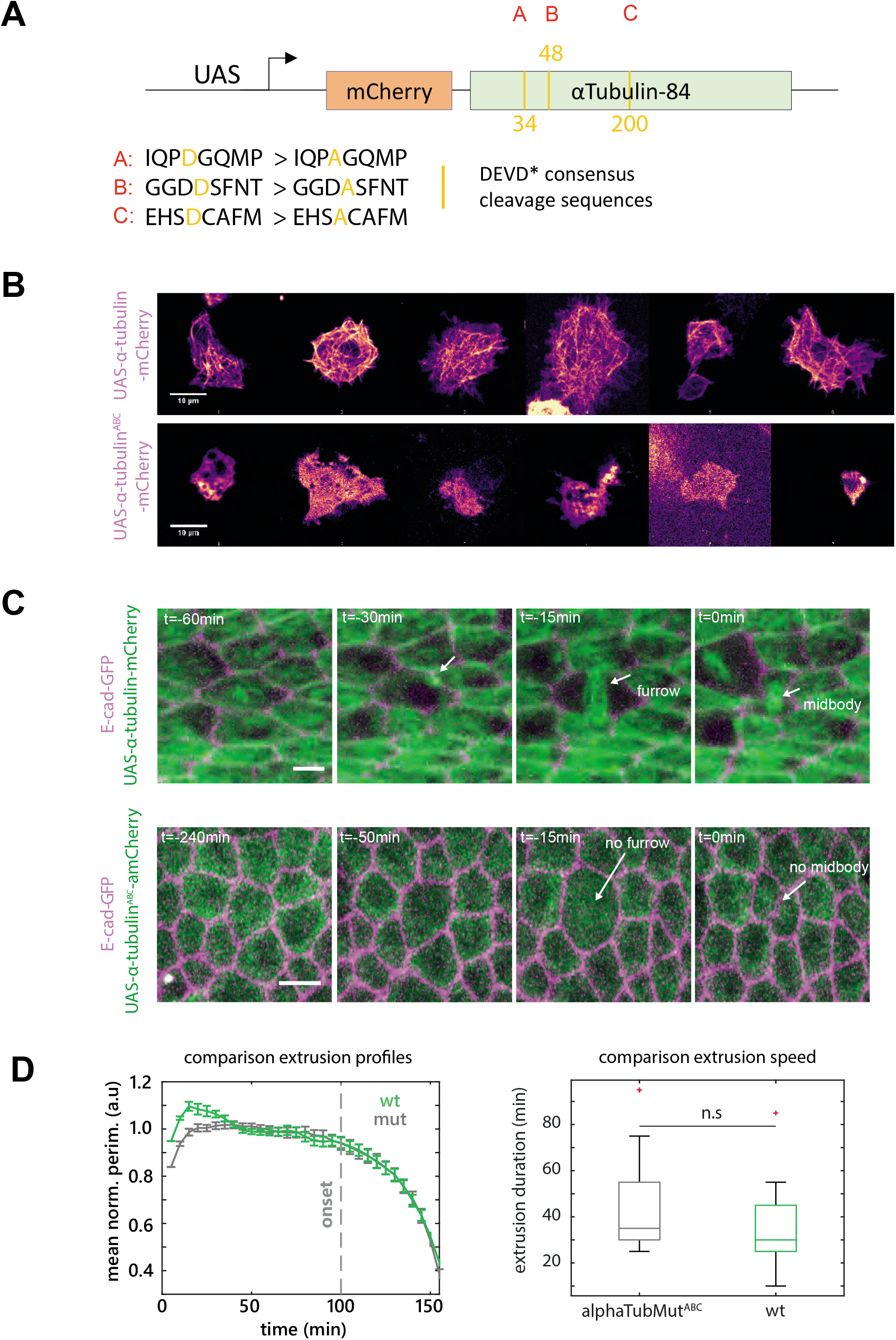
Mutation of αTubulin caspase cleavage sites prevents proper integration in MTs. **A:** Schematic representation of the α-Tubulin-mCherry construct. A,B,C shows the different caspase-cleavage sites at 34, 48 and 200 amino acids respectively. Yellow bars show mutation sites. The sequence of the cleavage site and the introduced mutations are shown below. **B:** Snapshots of different representative S2 cells transfected with the WT αTubulin-mCherry construction (top) or the mutated form (labelled α-Tubulin^ABC^-mCherry, bottom) driven by the UAS promoter. White arrows point at MTs in the WT form. These are absent from the cells expressing the mutant α-Tubulin^ABC^-mCherry (mostly cytoplasmic signal). **C:** Comparison of the α-Tubulin-mCherry and α-Tubulin^ABC^-mCherry localisation in the *Drosophila* pupal notum (UAS promoter, pnr-gal4 driver). Top row shows αTubulin-mCherry WT (green) with E-cad-GFP (magenta). White arrows point at one example of division furrow and the persisting midbody after cytokinesis. Bottom row shows mutant α-Tubulin^ABC^-mCherry. White arrows point at the absence of labelling at the division furrow or the midbody. Scale bars are 5μm. **D:** Left panel, Averaged and normalised apical perimeter of extruding cells expressing either WT UAS-α-Tubulin-mCherry (green) or mutant UAS-α-Tubulin^ABC^-mCherry (grey). Error bars are s.e.m.. Grey dotted line shows the onset of extrusion. Cells are aligned by the end of extrusion. n= 30 cells for both conditions. Right panel shows box blot of extrusion duration (between initiation and full apical closure) of cell expressing either UAS-α-Tubulin-mCherry (green) or UAS-α-Tubulin^ABC^-mCherry (grey). The difference is not significant (n.s.). n= 30 cells for both conditions.

## Supplementary movie legends

**Movie S1: E-cad evolution during cell extrusion**

Local projection of E-cad-GFP in an extruding cell from the midline region of the pupal notum. Scale bar is 5μm.

**Movie S2: MRLC evolution during cell extrusion**

Local projection of sqh-GFP (MRLC) in an extruding cell from the midline region of the pupal notum. Scale bar is 5μm.

**Movie S3: Actin evolution during cell extrusion**

Local projection of utABD-GFP in an extruding cell from the midline region of the pupal notum. Scale bar is 5μm.

**Movie S4: Localisation of Rho during cell extrusion**

Local projection of aniRBD-GFP (Rho localisation, left, green) and E-cad-tdTomato (right, magenta) in an extruding cell from the midline region of the pupal notum. Scale bar is 5μm.

**Movie S5: Vertex model based simulations of early steps of cell extrusion.**

**Left panel:** control simulation. The tracked cells (blue) have parameters values identical to all the other cells. **Middle panel:** purse-string driven extrusion. At t = 20 sts, the tracked cells (blue) were forced to initiate extrusion by increasing at each iteration their contractility parameter 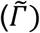 with a fixed rate 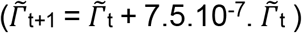. **Right panel:** At t = 20 sts, the tracked cells were forced to initiate extrusion by decreasing after each iteration their resting area (A_α_^(0)^) with a fixed rate (A_α_^(0)^t+1 = A_α_^(0)^t – 3.5.10^4^ A_α_^(0)^ t).

**Movie S6: Evolution of cell volume during extrusion**

3D rendering (top view left, lateral view, right) of an extruding cell upon activation of caspases by optoDronc. E-cad-tdTomato in red and cytoplasmic GFP in green. Scale bar is 5μm.

**Movie S7: Localisation of Talin during cell extrusion**

Local projection of Talin-GFP in an extruding cell from the midline region of the pupal notum. Scale bar is 5μm.

**Movie S8: Depletion of apical MTs in an extruding cell**

Local projection of Jupiter-GFP (total tubulin, green) and E-cad-tdTomato (magenta) in an extruding cell from the midline region of the pupal notum. Scale bar is 5μm.

**Movie S9: Depletion of EB1 in an extruding cell**

Local projection of EB1-GFP (MT plus end binding, green) and E-cad-tdTomato (magenta) in an extruding cell from the midline region of the pupal notum. Scale bar is 5μm.

**Movie S10: Depletion of human α-tub in an extruding cell**

Local projection of an extruding cell expressing human α-tub-mCherry (green) and E-cad-GFP (magenta) in the midline region of the pupal notum. Scale bar is 5μm.

**Movie S11: Dynamics of Shot in an extruding cell**

Local projection of Shot-GFP (green) and E-cad-tdTomato (magenta) in an extruding cell of the midline region of the pupal notum. Scale bar is 5μm.

**Movie S12: Dynamics of Patronin in an extruding cell**

Local projection of Patronin-GFP (green) and E-cad-tdTomato (magenta) in an extruding cell of the midline region of the pupal notum. Scale bar is 5μm.

**Movie S13: Evolution of MTs upon cell apical reduction during large scale ablations**

Square laser ablation in a pupal notae expressing E-cad-GFP (middle, magenta) and α-tub-mCherry (right, green). Scale bar is 25μm.

**Movie S14: MT depletion upon optoDronc activation with or without p35**

Local projection of sirTubulin (left), E-cad-tdTomato (middle) and GFP (expressed in optoDronc clones, right) upon caspase activation with optoDronc (top) or upon optoDronc activation combined with p35 expression (inhibitor of effector caspases, bottom). Scale bars is 10μm.

**Movie S15: Cell shape evolution upon local MT recovery**

Local projection of EB1-GFP (green, middle) and E-cad-tdTomato (magenta, right) in a colcemid injected pupae in a control region (top) or in a region exposed to repetive pulses of UV (bottom). Note the recovery of EB1 cometes and the increase of cell apical area. Scale bar is 10μm.

**Movie S16: Cell shape evolution upon partial MT depletion**

Local projection of sqh-mKatex3 (MRLC, green) and a clone expressing UAS-α-tub-GFP (magenta) and the clustering system LARIAT. Upon blue light exposure, GFP forms aggregate and cells tend to reduce their apical area. Scale bar is 10μm.

**Movie S17: Cell extrusion pattern upon MT depletion in clones by Spastin**

Live pupal nota expressing E-cad-GFP (green) with *UAS-RFP* clones (magenta, conditional induction) alone (left) or with UAS-Spastin (right). White dots show extruding cells outside the clone, orange dots extrusions inside the clones. Scale bar is 25μm.

**Movie S18: Cell extrusion pattern in hid-dsRNA clones upon MTs depletion**

Live pupal nota expressing E-cad-GFP (green) with *UAS-hid-dsRNA* clones (magenta) injected with water and ethanol (left) or colcemid (right). White dots show extruding cells outside the clone, orange dots extrusions inside the clones. Scale bar is 25μm.

**Movie S19: Examples of extrusion events with or without caspase activation**

Example of extruding cells from *UAS-hid-dsRNA* clones (marked with his-mIFP, 3^rd^ column, greyscale) in a pupae expressing E-cad-tdTomato (magenta, left), the effector caspase sensor UAS-GC3Ai in the clones (green, right), and injected with colcemid. Top: an extrusion without sign of GC3Ai activation, bottom, an extrusion positive for GC3Ai. Scale bar is 10μm.

**Movie S20: Cell constriction in optoDronc p35 upon colcemid injection**

Local projection of E-cad-tdTomato (middle, magenta) and GFP (expressed in optoDronc clones, right, green) upon Caspase9 activation with optoDronc while inhibiting effector caspases with p35. Top movie is a mock-injected pupae, bottom movie is a pupae injected with colcemid. Scale bars is 10μm.

